# Inhibitory neurons control the consolidation of neural assemblies via adaptation to selective stimuli

**DOI:** 10.1101/2023.04.25.538236

**Authors:** Raphaël Bergoin, Alessandro Torcini, Gustavo Deco, Mathias Quoy, Gorka Zamora-López

## Abstract

Brain circuits display modular architecture at different scales of organization. Such neural assemblies are typically associated to functional specialization but the mechanisms leading to their emergence and consolidation still remain elusive. In this paper we investigate the role of inhibition in structuring new neural assemblies driven by the entrainment to various inputs. In particular, we focus on the role of partially synchronized dynamics for the creation and maintenance of structural modules in neural circuits by considering a network of excitatory and inhibitory *θ*-neurons with plastic Hebbian synapses. The learning process consists of an entrainment to temporally alternating stimuli that are applied to separate regions of the network. This entrainment leads to the emergence of modular structures. Contrary to common practice in artificial neural networks – where the acquired weights are typically frozen after the learning session – we allow for synaptic adaptation even after the learning phase. We find that the presence of inhibitory neurons in the network is crucial for the emergence and the post-learning consolidation of the modular structures. Indeed networks made of purely excitatory neurons or of neurons not respecting Dale’s principle are unable to form or maintain the modular architecture induced by the entrained stimuli. We also demonstrate that the number of inhibitory neurons in the network is directly related to the maximal number of neural assemblies that can be consolidated, supporting the idea that inhibition has a direct impact on the memory capacity of the neural network.

## Introduction

Inhibition has an essential role for the dynamics and the adaptation ability of the brain^1,2^. Inhibitory interactions are mediated by the GABAergic neurotransmitters whose effect is to reduce the probability of their target neurons to emit spikes. This is usually true in the adult brain operating in normal conditions^3^. Inhibition has been shown to be fundamental for perceptual decision making^4^ as well for the emergence of brain rhythms^5^. It also plays a crucial role in adaptation processes of the brain, in order to shape and preserve the structure of the network according to the correlations between inputs^6–8^.

Many models have been proposed in the literature to study the properties of plastic neural networks that account for excitation and inhibition. For example, models that treat neurons as oscillators have been useful to explore the synchronization phenomena of the networks as a function of the ratio between excitation and inhibition^9,10^. Synaptic plasticity is usually mimicked by introducing adaptive coupling between the oscillators^11–13^, this can promote the emergence of multi-stable regimes characterized by synchronized and desynchronized clusters of oscillators^13–16^; analogous to observations in the presence of spike-timing-dependent plasticity (STDP)^17,18^.

While the simplicity of these models favours their analysis, they typically omit several biological aspects. Two major omissions are shared with common models for artificial neural networks employed in artificial intelligence and machine learning. On the one hand, excitation and inhibition are considered at the level of individual links and thus no distinction is made between excitatory and inhibitory neurons^11–13,19^. Despite the fact that from a biological point of view Dale’s principle^20^ requires that the nature of the neurons should be uniquely defined such that all post-synaptic connections of a neuron are either excitatory or inhibitory. On the other hand, plasticity is only accounted for during the learning phase in which the networks are entrained. After the training, the resulting synaptic weights are frozen. However, in biological neural networks adaptation is constantly active raising the question of how could memories be consolidated in a network that is susceptible to permanent change.

The aim of this paper is to study the role of inhibitory neurons in the emergence and consolidation of neural assemblies due to stimulus-driven plasticity. Therefore, we model networks of excitatory and inhibitory neurons (represented as *θ*-neurons^21^) shaped by external stimuli applied to differentiated subsets of neurons. The formation of assemblies is obtained thanks to a symmetric Hebbian-like phase-difference-dependent plasticity rule allowing the correlated neurons to reinforce their synaptic weights. Our results show that the presence of inhibitory neurons is crucial not only for the formation of clusters of neurons triggered by the stimuli, but also for the consolidation and the recall of memories. We find that violation of the biological constraints, either by constructing networks of excitatory-only neurons or by omitting Dale’s principle, leads to networks unable to maintain these memories. Furthermore, we show that the memory capacity of the network is directly related to the number of inhibitory neurons and that the conservation of the inhibitory synaptic weights is sufficient for memory recalls even if the synaptic patterns associated to the excitatory neurons are forgotten. The robustness of the results is tested against variations of the entrainment protocol and replacing the *θ*-neuron model by other models of oscillators such as the Kuramoto and the Stuart-Landau models.

## Background

Numerous studies have shown that the brain’s connectivity follows a modular organization at different spatial and functional scales, with neurons and regions associated to common modalities or functions being more strongly connected^22–26^. These modules are usually associated to particular sensory modality (e.g. vision, audition and motor control) or to specific features within a modality, emerging in an autonomous way^23,24,26–28^. Plastic connection strengths seem to play a significant role in the specification of neural assemblies involved in a particular function under the action of co-activation zones^29^. This highlights the concept of semantic memory where correlated information or functions share a common structure^30,31^. Some models suggest this concept of semantic memory by having association between mental representations and topology^32,33^. Also, it has been observed that during rest (no task activity) small series of sequence activation replays occur, akin to a memory retrieval and consequently to a process of memory consolidation^34,35^. This memory retrieval process in the dynamics seems based on the activation of particular semantic subgroups^34^, again highlighting the impact of the physical organization of the network on these dynamics.

Regarding inhibition, it has been proposed that the interconnections between excitatory and inhibitory cells play a relevant role balancing between fast adaptation and long-term conservation of the memories in neural circuits^36,37^. The problem of consolidating over the long term the memories a network has learned, while keeping an on-going adaptation process that avoids forgetting past memories by new ones, is a relevant issue encountered in models of various fields from biology to machine learning and artificial intelligence. In the brain, the connections between excitatory neurons are more volatile implying both forgetting in the absence of inputs and relearning in the presence of new information^36^. Therefore, it seems that while excitatory plasticity initially shapes the connectivity, driven by the input information^36,38–40^, long-term maintenance appears to be mediated by the inhibitory plasticity^36^. Indeed, it has been shown that inhibitory plasticity has a fundamental role in balancing excitation and inhibition in recurrent networks, as well as in storing synaptic memories^41^. Inhibitory neurons can modulate the level of synchronization of the network, making it possible to preserve consistency in the dynamics of modular areas^10^. The interplay between excitatory and inhibitory currents also makes it possible to have a more efficient coding^42,43^, which can be of great interest especially in the field of artificial intelligence.

## Methods

This section describes the model employed for the dynamics of the membrane potentials of the neurons in the network, the learning rules governing the weight adaptation and the protocols followed to induce structural patterns and their consolidation, as well as the protocols used to analyse the stability of the emerged patterns.

### Neuronal dynamics

To mimic the membrane potential evolution of a neuron we employ the *θ*-neuron model introduced by Ermentrout and Kopell as the normal form for class I excitable cells^21^. This model has the advantage of reproducing quite faithfully the behaviour of spiking neurons through the description of the evolution of the corresponding “membrane potential” while remaining a phase model^44^. Indeed, it can be put in an one to one correspondence with the Quadratic Integrate and Fire (QIF) spiking neuronal model via a non-linear transformation linking the phase to the membrane potential^45^. Furthermore, the *θ* model can describe bursting dynamics observable at the level of the membrane potential of specific cells of the Aplysia mollusc^21,44^. In addition, networks of coupled *θ*-neurons can exhibit a variety of behaviours ranging from asynchronous regimes to multi-stability, from partial synchronization to chaos^46–49^. In a network of *N* = *N*_*E*_ + *N*_*I*_ neurons, where *N*_*E*_ (*N*_*I*_) is the number of excitatory (inhibitory) neurons, the dynamical evolution of the phase *θ*_*i*_ of a neuron (*i* = 1, …, *N*) is governed by the following equation:

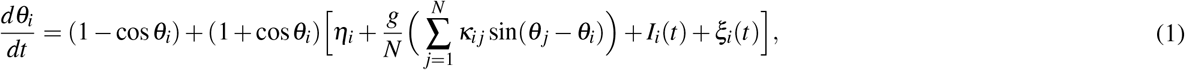

where *η*_*i*_ is the excitability or the bifurcation parameter controlling the intrinsic frequency of the neuron, *g* represents a global synaptic strength modulating the overall level of coupling among the neurons, and *κ*_*i j*_ is a matrix term measuring the relative synaptic weight from the pre-synaptic neuron *j* to the post-synaptic neuron *i*. The latter term is subject to a temporal evolution, therefore we can speak of adaptive coupling, the evolution rules for the terms *κ*_*i j*_ will be introduced in the next sub-section. Furthermore, we consider a diffusive sinusoidal coupling function based on the phase differences as in the Kuramoto model^50^. In other words, the couplings between neurons are considered as electrical gap junctions. The phases *θ*_*i*_ are defined in the range [−*π, π*[and we assume that whenever the phase *θ*_*i*_ reaches the threshold *π* a spike is emitted by the *i*-th neuron.

External inputs are incorporated via the current terms *I*_*i*_(*t*) representing time-dependent stimuli. Positive external currents increase the frequency of the neurons, thus imitating the typical biological response of sensory neurons exposed to stimuli of varying intensity. Finally, an independent Gaussian noise *ξ*_*i*_(*t*) term is applied to each neuron mimicking background noise always present in real neural circuits.

Simulations are performed by integrating Eq. (1) via an Euler scheme, where the multiplicative stochastic term is treated in the Stratonovich sense^51^, with an integration time step *dt* = 0.01.

In order to quantify the degree of synchronization in the network, we introduce the Kuramoto-Daido order parameters *Z*_*n*_(*t*)^52–54^. The *n*^*th*^ order parameter is defined as follows:

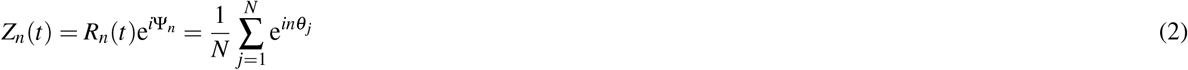

where *R*_*n*_ is the modulus of the complex order parameter *Z*_*n*_ and Ψ_*n*_ the corresponding phase. The modulus of *Z*_1_ is employed to characterize the level of phase synchronization in the network: *R*_1_ *>* 0 (*R*_1_ = 1) for a partially (fully) synchronized network, while 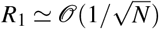 for an asynchronous dynamics. The other modulus *R*_*k*_ with *k >* 1 are used to characterize the emergence of multi-clusters, in particular *R*_2_ is finite whenever two clusters in anti-phase are present in the network.

### Learning and adaptation

#### Plasticity function

The synaptic weights *κ*_*i j*_ are modified according to a symmetric Hebbian-like rule depending on the instantaneous phase differences between the neurons instead of spike timings as observed in real neurons^55^. Since *θ*-neurons are phase oscillators this choice allows to have a relatively precise temporal adaptation – as compared to a frequency based rule – that can be numerically integrated simultaneously with the neuronal dynamics. A symmetric plasticity facilitates the convergence of the weights and highlights better the correlations with respect to the inputs. Nevertheless it shall be noted that a symmetric plasticity is possible in this case because the transmission delays are ignored^56–58^.

We first consider the plasticity function introduced by Aoki et al.^11,12^ and recently employed in Berner et al.^13,16^. Given the phase difference Δ*θ* = (*θ*_*j*_ −*θ*_*i*_), this plasticity function is defined as:

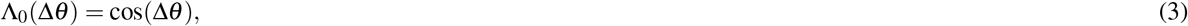

see Fig. 1, blue line. This rule follows the principle of Hebbian learning with a reinforcement (depression) of the weights when the neurons are in phase (in anti-phase). However although simple, the relevance in terms of realism of this function can be questioned due to the symmetry of the positive and negative parts, corresponding to long-term potentiation (LTP) and depression (LTD), respectively. Indeed, in biology LTP and LTD do not act on symmetric time windows^59,60^. To remedy this, we propose a plasticity function inspired by Lucken et al.^61^ and Shamsi et al.^62^. We define the new plasticity function as:

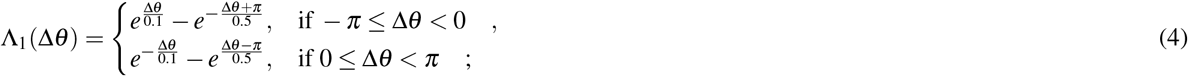

**Figure 1.**
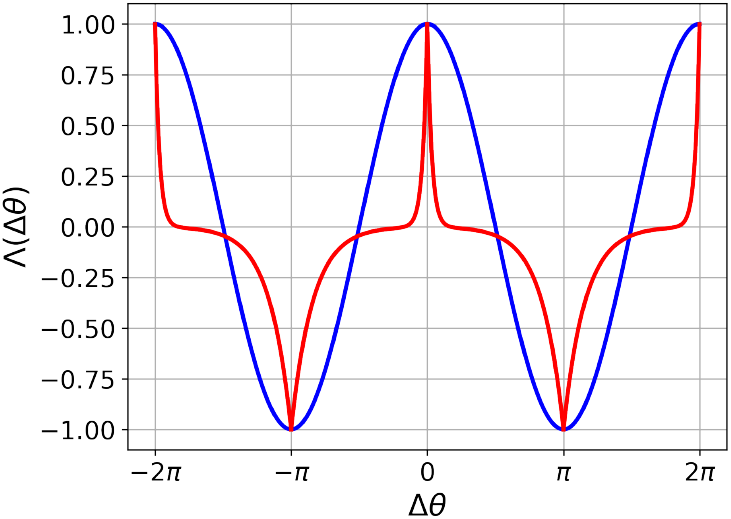
Hebbian phase difference-dependent plasticity functions Λ(Δ*θ*) versus Δ*θ*. In blue the function Λ_0_ reported in Eq. (3) and in red the function Λ_1_ defined Eq. (4). Despite having different potentiation and depression phase windows, the two functions attain the same maximum and minimum values.

where we assume that Λ_1_(Δ*θ*) is 2*π* periodic as shown in Fig. 1, red curve. This rule follows the same general principle of Hebbian learning as the one in Eq. (3) but the cosine is replaced by a combination of exponential functions making the potentiation (where Λ_1_(Δ*θ*) *>* 0) and the depression (where Λ_1_(Δ*θ*) *<* 0) phase intervals asymmetric. Since for phase oscillators, the phase evolution can be mapped into a time evolution, these asymmetric phase intervals correspond to asymmetric time windows. Indeed, this asymmetry is more consistent with the biological features: as a matter of fact, in the cortex depression operates over a longer time window with respect to potentiation^63,64^. Specifically in^60^, the authors have shown that for CA3 pyramidal cells in rat hippocampus, the STDP presents a depression time scale five times longer than the potentation one, analogously to our choice in Eq. (4).

In the following, unless stated otherwise, we will employ the function Λ_1_ and for simplicity we will refer to it as Λ(Δ*θ*).

#### Slow and fast adaptation

External stimulations have a direct impact on the adaptation of the synaptic weights. In presence of stimulation the synaptic weights should be modified quite fast to retain the information contained in the stimuli. However, in the absence of further stimuli, a fast adaptation may allow for a rapid relaxation of the weights towards their equilibrium values leading to a loss of the learned input-driven memories. This problem is common to most learning neural network models. The solution typically adopted consists in freezing the weights once the stimuli have been delivered. There are indications that two learning processes may occur simultaneously or sequentially in the brain. An example, is represented by the complementary mechanisms involved in short-term or long-term depression and potentiation of the synapses^64–67^. There are also evidences of different learning schemes on proximal and distal dendrites of pyramidal neurons^68^ or in the retina^69^. Finally, it is also worth to mention that long-term potentiation may be modulated by homeostatic plasticity^70,71^. Two learning processes also occur when acquiring new skills: a fast learning in a cortical structure is simultaneously slowly learned in a subcortical structure (habit learning for instance)^72–75^. Thus, according to previous experiences and the nature of inputs, the rate of learning may need to evolve so that neurons adapt more or less quickly to sensory inputs^69,72,76,77^. More precisely, adaptation in the brain is the result of multiple learning processes acting at distinct time scales allowing more or less rapid learning or relearning of new information without forgetting older memories^39,72^. To somehow mimic this phenomenon, we introduce two complementary learning time scales:

- a fast time-scale that activates in the presence of an external stimulus presented to the pre-synaptic neuron;
- a slow steady time-scale that has a minimal short-term effect.

The fast and slow time scales have been introduced in this phase model, where adaptation depends on the phase differences and not on the spiking times, to reproduce a fast learning period induced by the higher firing rate of the stimulated neurons and a slower consolidation phase, where the neuronal firing becomes sparse.

Given the plasticity function in Eq. (4) and the constraints just mentioned, we define the evolution of the coupling weights |*κ*_*i j*_|≤ 1 from pre-synaptic neuron *j* to post-synaptic neuron *i* to be governed by the following equation:

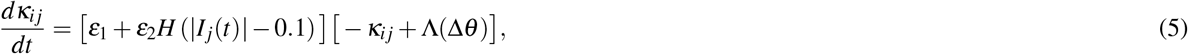

where *I*_*j*_(*t*) is an external input to neuron *j, H*(*x*) is the Heaviside function such that *H*(*x*) = 1 if *x >* 0 and *H*(*x*) = 0 otherwise, and *ε*_1_ *« ε*_2_ *«* 1 are the learning rates for the slow and the fast adaptations, respectively. The Heaviside function has a thresholding effect allowing fast adaptation only if a sufficiently strong stimulus |*I*_*j*_(*t*)| *>* 0.1 is presented.

#### Excitatory and inhibitory neurons

In order to account for the excitatory and inhibitory nature of the neurons, we disentangle Eq. (5) into two different cases, where the synaptic weights are bounded between [0, 1] ([− 1, 0]) provided the pre-synaptic neuron is excitatory (inhibitory). The dynamics of the weights are thus governed by the following equations.

If both neurons *i* and *j* are excitatory, then:

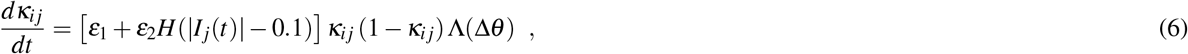

if the pre- and/or post-synaptic neuron is inhibitory the weight evolution is given by:

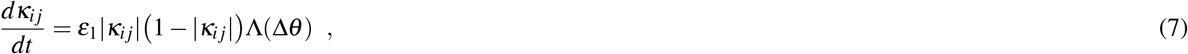

where 0 ≤ *κ*_*i j*_ ≤ 1 (−1 ≤ *κ*_*i j*_ ≤ 0) if the pre-synaptic neuron is excitatory (inhibitory). The non-linear dependence on *κ*_*i j*_ introduced in Eqs. (6) and (7) is intended to mimic the soft bounds often employed in the implementation of the STDP in spiking neural networks^78^ and to maintain |*κ*_*i j*_| smaller than one.

Notice that the fast adaptation is only present for synapses connecting excitatory neurons since external inputs are only applied to this type of neurons^79–82^. Indeed, sensory inputs must be rapidly assimilated in the short-term while being volatile in order to learn new information^36,72,83^. Therefore, synaptic weights involving inhibitory neurons will always only evolve on the long time scale. By default, we consider a ratio of 80% of excitatory neurons and 20% of inhibitory neurons as is commonly accepted in the human cortex^7,10,36^.

Lastly, we expect that for synapses connecting two excitatory or inhibitory neurons *i* and *j*, the weights *κ*_*i j*_ and *κ*_*ji*_ will become equal in the long run, due to the symmetry of Eq. (6) in the absence of stimulation and of Eq. (7). However, this should not be the case if one of the neurons is inhibitory and the other excitatory.

### Stimulation protocols

#### Main experiments

We now summarise the principal protocol followed in the paper. The networks are initialised with random weights uniformly distributed in the range *κ*_*i j*_ ∈ [0, 1] when the neuron *j* is excitatory and in the range *κ*_*i j*_ ∈ [−1, 0] when the neuron *j* is inhibitory. We consider heterogeneous neurons with excitabilities *η*_*i*_ randomly distributed according to a normal distribution 𝒩 (1.5, 0.01). The initial phases of the neurons are randomly selected in the interval [−*π, π*]. All network parameters are summarized in Table 1.

**Table 1.**
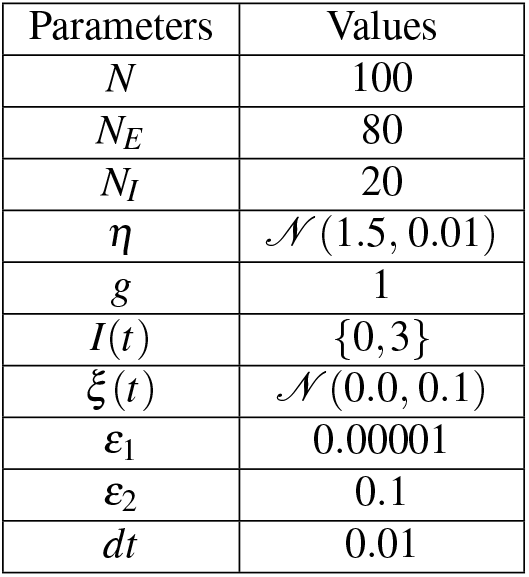
Parameters for the network of *θ*-neurons

| Parameters | Values |
| --- | --- |
| $N$ | 100 |
| $N_E$ | 80 |
| $N_I$ | 20 |
| $\eta$ | $\mathcal{N}(1.5, 0.01)$ |
| $g$ | 1 |
| $I(t)$ | $\{0, 3\}$ |
| $\xi(t)$ | $\mathcal{N}(0.0, 0.1)$ |
| $\varepsilon_1$ | 0.00001 |
| $\varepsilon_2$ | 0.1 |
| $dt$ | 0.01 |

The stimulation protocol is illustrated in Fig. 2. Initially, a period of spontaneous activity is considered in order to allow the system to relax to its rest state. Afterwards, a learning phase follows during which two external positive currents are applied randomly and intermittently for constant periods of 20 time units. One current stimulates the first half of the excitatory neurons (*i* = 1, 2, …, 40), and the other one the second half (*i* = 41, 42,…, 80). Inhibitory neurons are never directly stimulated. Finally, after the learning phase the network is left to evolve spontaneously for a long time. During this post-learning phase no neurons are stimulated and therefore only the slow adaptation rate *ε*_1_ remains active, affecting the stabilization of the connectivity patterns formed during the stimulation phase.

**Figure 2.**
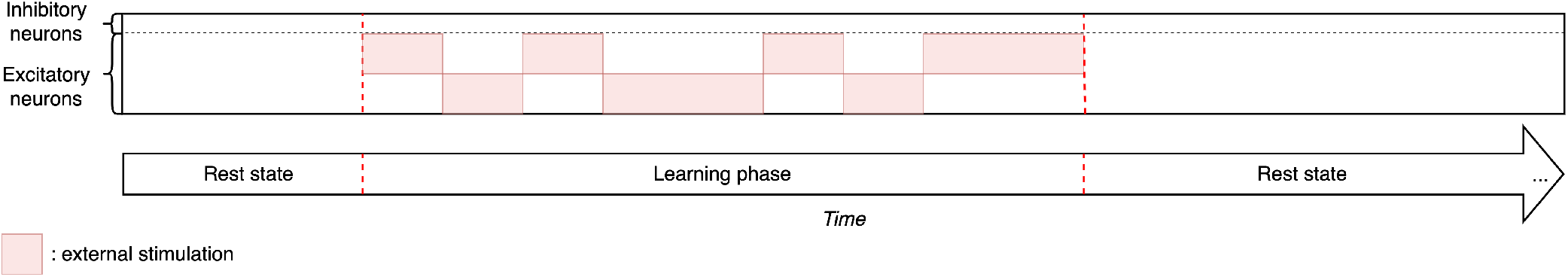
Diagram representing the stimulation protocol divided in three phases: a first short phase of spontaneous activity, a learning phase during which two different groups of excitatory neurons are stimulated alternatively over time (the red areas represent the duration and the neurons that received external stimulation), and a final long period of resting-state activity in which the plasticity remains active.

This principal experiment was repeated for different variations. In one case, we ignored inhibition and considered networks of only excitatory neurons. In a second case, in order to show the relevance of Dale’s principle for pattern consolidation, i.e. the need to preserve the nature of the excitatory and inhibitory neurons during the simulation, we considered unlabelled neurons. In this case neurons were allowed to display both excitatory and inhibitory synapses. Specifically, for the experiment with unlabelled neurons the synaptic adaptation was ruled by Eq. (5) with the Hebbian function Λ_0_(Δ*θ*) in Eq. (3).

#### Additional experiments

Thereafter variations of the principal protocol are considered, as follows:

1. three external stimuli during the learning phase instead of two;
2. an overlap of 8 neurons among two groups of excitatory neurons stimulated by two external stimuli;
3. a recall experiment of previously stored items, where the trace of the memory storage is maintained only in the connections involving inhibitory neurons.

Other variations of the main protocol are described in section *Alternative Stimulation Protocols* of the Supplementary Information.

To understand the generality of the results obtained in the additional experiments (1) and (2), we analyse the stability of different configurations. In particular, we perform a stability analysis where the objective is to determine the conditions under which the network connectivity during the resting state remains stable (i.e. the learned modular structure is preserved) or unstable (i.e. at least one structural cluster is not maintained) over the long term. This stability analysis corresponds in experiment (1) to the maintenance of all the stored clusters and in experiment (2) to the maintenance of the two stored clusters despite their increasing overlap.

In order to perform this analysis, we start from already trained networks satisfying the desired constraints (i.e. the number of structural clusters or the number of overlapping neurons between clusters versus the number of inhibitory neurons), thus facilitating the replication of experiments under different conditions. Moreover, this pre-training facilitates the analysis of extreme cases where for instance a single inhibitory neuron is associated with each cluster whose emergence is not necessary guaranteed via a random learning. Then we leave the network in a state of spontaneous activity for a long time and compare the final structural state to which it converges with the initially stored one to assess the stability or instability of this latter configuration.

## Results

The goal of this paper is to study the emergence and consolidation of modular architectures induced via a learning process promoted by localised inputs which target distinct subsets of neurons. We will first present results concerning the emergence and consolidation of clustered architecture induced by learning of two external stimuli. This experiment is repeated for networks composed of different types of neurons: namely, purely excitatory neurons, excitatory and inhibitory neurons, and unlabelled neurons. Then, we will study variations of the initial protocol and in particular the learning process promoted (i) by multiple stimuli and (ii) by overlapping stimuli, where neurons can encode for multiple items. Lastly, we investigate the ability of the neural networks to recall the stored memories.

### Learning and consolidation of modular assemblies

We start by investigating the emergence of two modules due to stimulations of non-overlapping neural populations. We first study this phenomenon in networks containing excitatory and inhibitory neurons and then we explore cases with only excitatory neurons or with unlabelled neurons whose connections can be either excitatory or inhibitory.

### Networks of excitatory and inhibitory neurons

We consider a network of *N* = 100 *θ*-neurons subdivided in *N*_*E*_ = 80 excitatory and *N*_*I*_ = 20 inhibitory ones, initially connected via a random weighted matrix (uniform distribution between [0, 1] or [−1, 0] for pre-synaptic excitatory or inhibitory neurons, respectively), see *Methods*. The network is entrained using two independent stimuli applied to separate subsets of neurons following the protocol shown in Fig. 3a. The phases of the neurons are randomly initialised (uniform distribution between [−*π, π*]). The results of this experiment are reported in Fig. 3b. During the initial rest phase the network is left to evolve spontaneously and it converges into a state close to synchrony despite the fact that the weight matrices at time T0 are randomly distributed, see raster plot (1). Both excitatory and inhibitory neurons tend to synchronize, as shown in the raster plot (2) at T0. This is due to the value of the chosen coupling strength *g*, which is sufficiently large to favour a synchronized phase.

**Figure 3.**
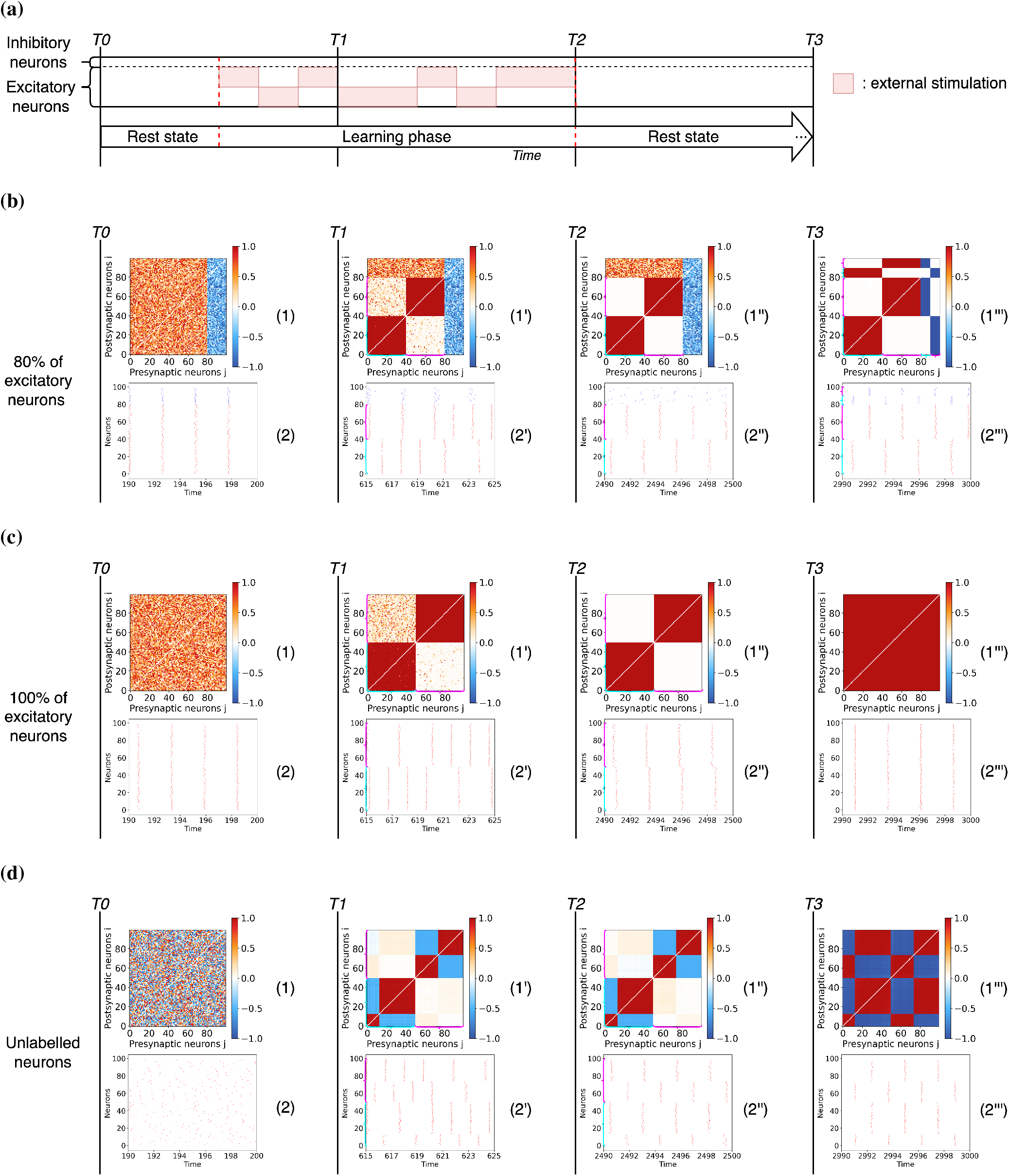
Entrainment of networks of *θ*-neurons defined by different combinations of excitatory and inhibitory connections. (a) Experimental protocol consisting of the stimulation of two non-overlapping neuronal populations of *θ*-neurons with plastic synapses. Stimuli are presented in temporal alternation. (b) Results for a network with 80% excitatory and 20% inhibitory neurons. (c) Results for a network of only excitatory neurons and (d) for unlabelled neurons projecting both excitatory and inhibitory postsynaptic connections. The results are reported at different moments of the stimulation. The time T0 corresponds to the beginning of the simulation before the learning phase, after a transient period *t*_*t*_ = 190 has been discarded. The time T1 corresponds to an early moment during the learning phase. Time T2 corresponds to the end of the learning phase and the beginning of the resting-state. The time T3 corresponds to the end of a long period of spontaneous activity during which synaptic weights are consolidated. Panels labelled (1), (1’), (1”) and (1”‘) represent the weight matrices at times T0, T1, T2 and T3: the color denotes if the connection is excitatory (red) or inhibitory (blue) or absent (white). Panels (2), (2’), (2”) and (2”‘) are raster plots at times T0, T1, T2 and T3, displaying the firing times of excitatory (red dots) and inhibitory (blue dots) neurons. Note that the inhibitory neurons are sorted by phases at time T3 in the weight matrices and raster plots reported in (b) and (c); while the neurons are sorted by phase in the weight matrices and raster plots within each cluster at times T1, T2 and T3 of the row (d). The cyan and magenta brackets represent clusters 1 and 2 respectively when they are visible in weight matrices and raster plots.

After the period of spontaneous activity the learning phase starts. The resulting connectivity matrix and the network dynamics are shown at two intervals: during the learning phase – time T1 – and at the end of this phase – time T2. As we see, the presence of the two stimuli leads to the emergence of two modular structures (weight matrices (1’) and (1”)) among the excitatory neurons while the weights involving inhibitory neurons do not evolve much due to the separation of fast and slow learning rates. The presence of an input leads to an increase in the firing rate of the stimulated neurons, see raster plot (2’). At the end of the learning phase, two disconnected clusters of excitatory neurons firing in anti-phase emerge in the network, while the inhibitory neurons remains uncorrelated (see raster plot (2”)).

Following the learning period the network is left to evolve driven by its spontaneous activity. In this stage the adaptation is governed by the slow adaptation rate. The long-term results are shown in the time column T3. As seen, the learned structure is consolidated. The two excitatory clusters are maintained, see weight matrix (1”‘), and besides this the input/output inhibitory weights continue adapting such that the modular structure is reinforced. Finally, also the inhibitory neurons split in two structural clusters. The final consolidated connectivity structure is reported in Fig. 4a. From this figure it emerges that the neurons are organised in two clusters, each one involving a group of excitatory neurons that projects in a feedforward manner on a group of inhibitory neurons. Therefore the inhibitory neurons within the first cluster synchronise with the excitatory neurons driving them, as shown in panel (2”‘). Furthermore, the inhibitory neurons of one cluster projects on the excitatory and inhibitory neurons of the other cluster. This induces a repulsion of the dynamics of the two clusters that adjust in anti-phase one with respect to the other, both clusters displaying exactly the same period of oscillation, see raster plot in panel (2”‘) in Fig. 3b. In order to validate the robustness of the results, the experiment was repeated several times (as all the following experiments) by randomly varying the presentation order of the stimuli to the two populations. At the same time, the initial values of the phases of the neurons and of the connection weights were also randomly assigned in each realization. From these data we have estimated the evolution in time of the modulus of the 1^*st*^ and 2^*nd*^ Kuramoto-Daido order parameters (as defined in Eq. 2). These indicators are useful to analyse the level of synchronization in the network and to test for the emergence of clusters at different stages during the experiment. The temporal evolution of the average order parameters and their standard deviations are shown in Fig. 4b for the previous experiment displayed in Fig. 3b.

**Figure 4.**
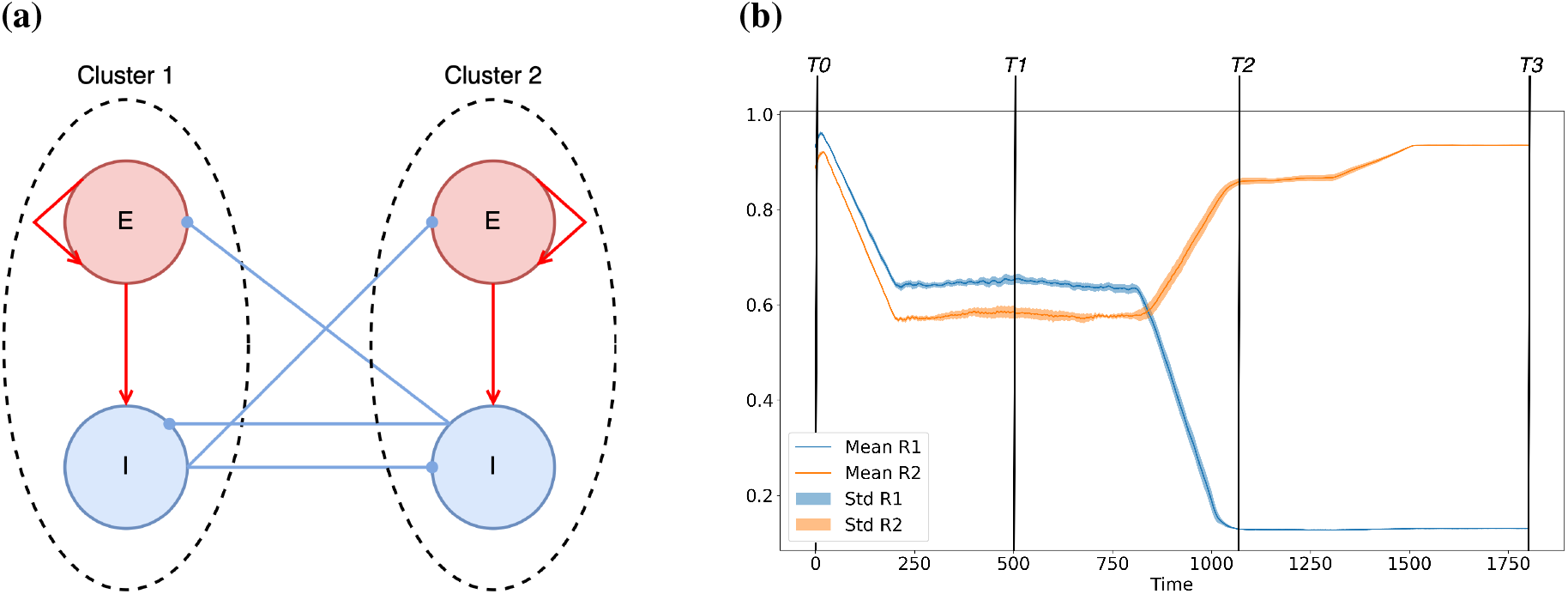
(a) Schematic diagram representing the connectivity matrix emerging after the consolidation phase for the stimulation of two non overlapping populations reported in Fig. 3. Red (blue) circles represent excitatory (inhibitory) populations, dashed circles identify clusters of synchronized neurons. (b) Evolution of the modulus of the 1^*st*^ (in blue) and 2^*nd*^ (in orange) Kuramoto-Daido order parameters Eq. (2) averaged over 10 different realizations randomised initial conditions and stimulation sequences. The mean and the corresponding standard deviation are shown versus time during the realization of the experiment in Fig. 3. The time labels have the same meaning as in Fig. 3.

At time T0, we observe with a negligible variation that the network is almost totally synchronized, as testified by the fact that *R*_1_ ≃1 and *R*_2_ ≃ 1. During the learning phase (i.e. at time T1), the value of the two parameters drop and their standard deviations increase, thus testifying for a larger variability and a desynchronization of the network. After the learning at time T2, *R*_2_ (*R*_1_) increases (decreases) tending toward a large (small) value. These results indicate the emergence of two main synchronized clusters in the network corresponding to the two learned modules. Finally, after the long term consolidation (i.e. at time T3), the network converges towards two clusters in anti-phase where *R*_2_ ≃ 1 (*R*_1_ ≃ 0) and the corresponding standard deviations are negligible, thus confirming the stability of this final state, despite the different stimulation sequences employed in the performed experiments. These observations put in evidence that the formation and consolidation of input-driven modules are a robust outcome of the model, which depends mainly on the characteristics of the stimulated populations and on the number of inhibitory neurons in the network, and is not significantly affected by the variability of the initial conditions nor of the particular sequence of the presented stimuli.

### Networks of excitatory neurons

Now we consider the case in which all neurons are excitatory and no inhibition is present. As before, the neurons synchronise during the initial stage of spontaneous activity and the training phase leads to the formation of two structural clusters (see, Fig. 3c). However, from the dynamical perspective the neurons in the two clusters are no longer firing in anti-phase, as in the previous case. Instead, at the end of the learning phase they present a small phase-shift and they indeed tend to synchronize as evidenced from the raster plot at time T2 (2”). As a consequence, during the stabilization phase, all the synapses are reinforced leading to an all-to-all connectivity matrix (see matrix (1”‘) at time T3). Therefore the two structural modules that emerged during the learning are now completely forgotten.

### Networks of unlabelled neurons

As a last case, we consider the typical scenario of artificial neural networks, where neurons are unlabelled, meaning that each neuron can display both excitatory and inhibitory post-synaptic connections violating Dale’s principle. The results of this experiment are reported in Fig. 3d. This time the initial stage of spontaneous activity leads to an asynchronous neuronal state (raster plot (2)) since every neuron randomly attracts and repulses its neighbours via excitatory and inhibitory links. After the learning phase (time T2), two symmetric structural clusters emerge in the weight matrix (1”). More precisely, the Hebbian rule in Eq. (3) leads to the creation within each structural module of two phase clusters in anti-phase, depending on the initial phases as explained in Aoki et al.^11,12^.

In contrast with the previous cases, the structural modules are now not well defined. Weak connections among the two clusters are still present as shown in (1”), allowing the possibility for the two clusters to synchronize, see the spike trains in (2”). During the consolidation phase these weak cross-modular connections increase, leading to a fully coupled matrix with no structural modules, see (1”‘). However, even in the absence of structural modules one can observe clustered phases in the temporal evolution of the neurons as also reported in^13^. The neurons connected through excitatory (inhibitory) pre-synaptic links tend to fire in phase (anti-phase). Nevertheless, these phase clusters do not reflect the two structural modules previously stored in the network.

In summary, we have shown that the presence of both excitatory and inhibitory neurons is needed for the formation and the consolidation of structural modules driven by the learning of independent stimuli. In the following, we generalize these results to cases in which the number of stimuli – and therefore of expected modules – is arbitrary and to the case where the stimuli act on overlapping neuronal populations.

### Learning multiple clusters

In this experiment, the previous learning protocol is repeated for networks with excitatory and inhibitory neurons but considering now three inputs as illustrated in Fig. 5a. The results are analogous to the previous ones, apart that in this case three clusters emerge. As before during the final consolidation stage each inhibitory neuron ends associated to one of these clusters, connectivity matrix (1”‘) in Fig. 5b. From the dynamical point of view, the neurons of the different clusters no longer fire in anti-phase, but instead at regular intervals corresponding to one-third of the firing period of the single cluster, see the raster plot (2”‘) in Fig. 5b.

**Figure 5.**
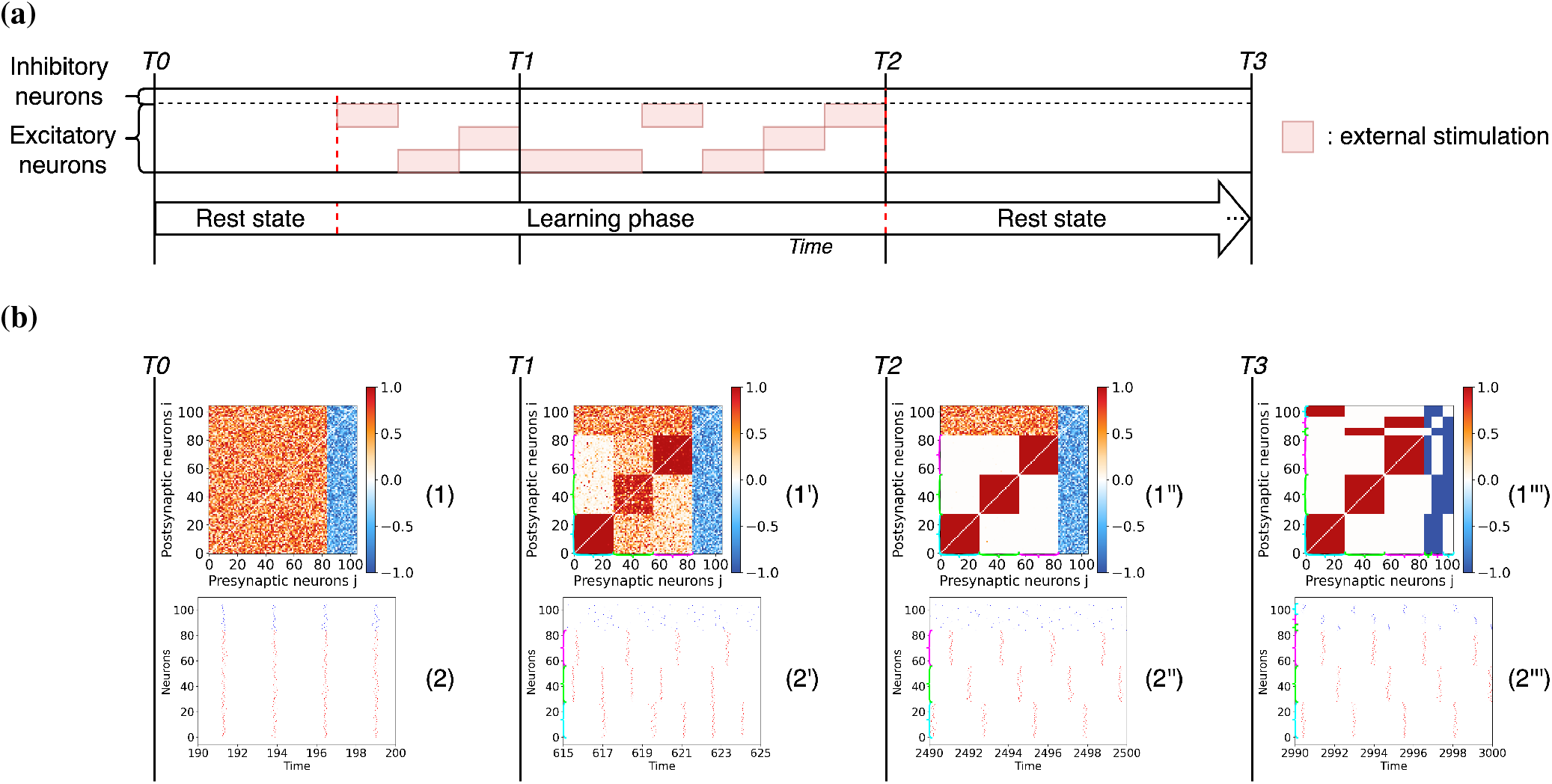
Entrainment of a networks of excitatory and inhibitory *θ*-neurons with three stimuli. (a) Stimulation protocol of the excitatory neurons by three different and non-overlapping stimuli. (b) Entrainment results at different instances of the simulation. The time labels and the graphs have the same significance and content as in Fig. 3. Inhibitory neurons are sorted by phases at time T3 for visualization purposes. The cyan, green and magenta brackets represent clusters 1, 2 and 3 respectively when they are visible in weight matrices and raster plots.

These results generalize by increasing the number of independent inputs. However, when there are too many inputs, we observe that it is more difficult for the network to sustain separate dynamics for each cluster and therefore the long-term consolidation is compromised. The reason is that, as the number of inputs increases, the clusters are made of fewer neurons and it becomes less likely that a sufficient number of inhibitory neurons will associate with each cluster allowing for its consolidation. This opens up the question about how many inhibitory neurons are necessary in the network in order to maintain clusters that fire at distinct times.

To answer this question, we perform the following experiment: we artificially prepare weight matrices made of *M* modules of excitatory neurons similar to those found at the post-learning time T2. Then, we associate each available inhibitory neuron to a single structural module such that it receives excitatory inputs from this cluster and inhibits the excitatory neurons of all the other clusters. Thus generalizing the architecture reported in Fig. 4a to many clusters, each containing a single inhibitory neuron. Starting from these weight matrices, we let the network spontaneously evolve for a long period, similarly to what was previously done during the post-training phase, and at the end of the simulation we count the number of structural modules that survive. We consider *N* = 100 neurons in the network and repeat the experiment for an increasing number *M* of modules (namely, from *M* = 2 to *M* = 100) controlled by varying the number of inhibitory neurons.

The results are shown in Fig. 6a. We find that in order to maintain *M* independent modules the network needs to contain at least *M* − 1 inhibitory neurons. An example of the effect of this limitation is shown in Supplementary Figure 4, where the experiment of Fig. 5 is repeated for a network containing a single inhibitory neuron. In this case even if the original weight matrices contained three structural clusters only two independent clusters could be maintained in the end.

**Figure 6.**
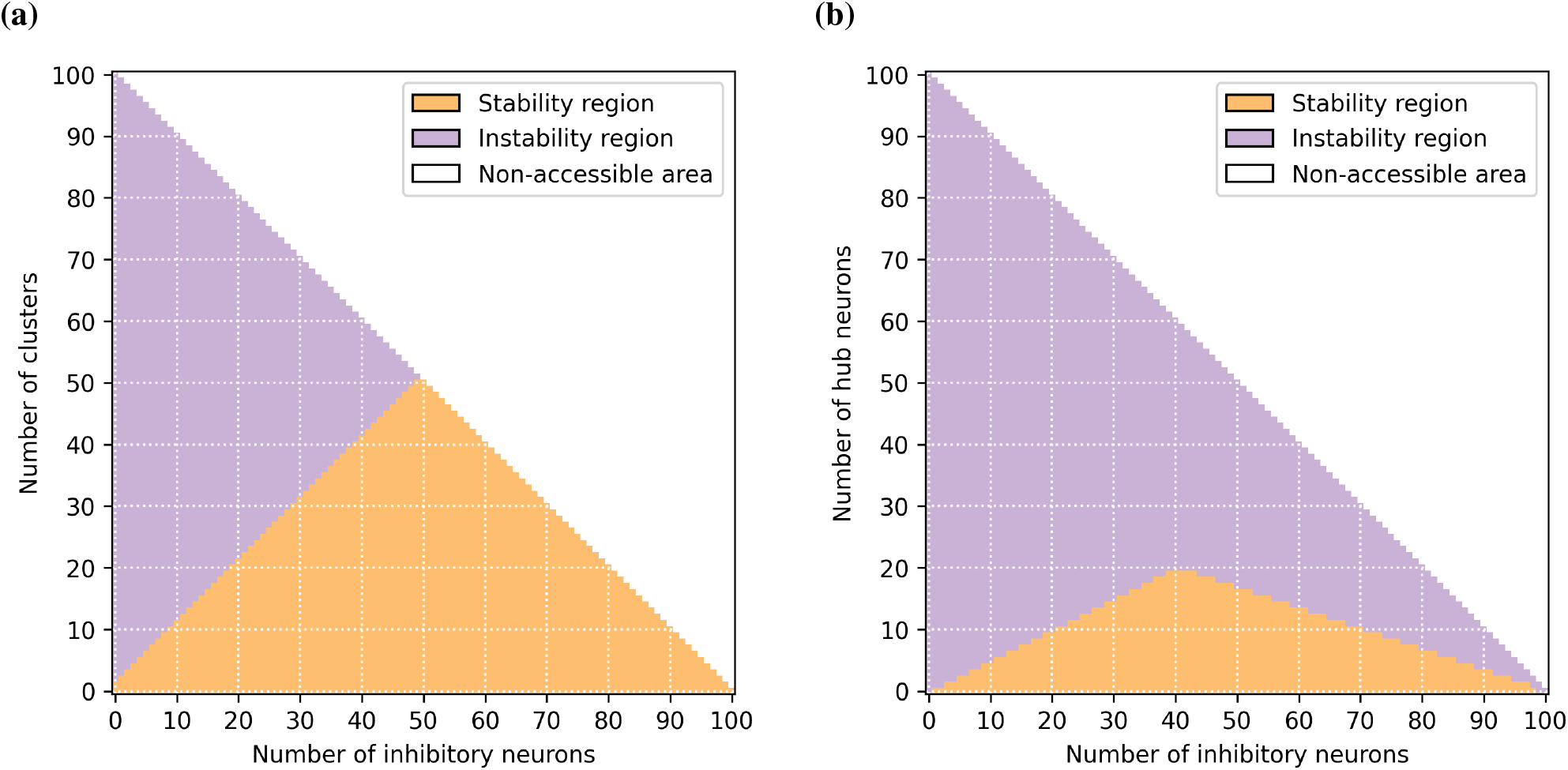
(a) Regions of stability (orange) and instability (purple) of the structural modules versus the number of clusters initially present in the weight matrix and the number of inhibitory neurons in the network. The line separating the two regions represents the upper limit to observe *M* independent clusters versus the number of inhibitory neurons. This limit corresponds to *M* −1 inhibitory neurons. (b) Regions of stability (orange) and instability (purple) of the two clusters versus the number of excitatory neurons encoding for the two stimuli (hub neurons) and the number of inhibitory neurons in the network. The lines separating the two regions represent the minimal number of inhibitory neurons *N*_*I*_ required to observe two clusters with *N*_*H*_ hubs. For 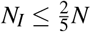 the line is given by *N*_*I*_ = 2*N*_*H*_ + 1. For larger number of inhibitory neurons the limit is given by *N*_*I*_ = *N* −3*N*_*H*_. Note that both graphs have been realized for a network made of *N* = *N*_*E*_ + *N*_*I*_ = 100 neurons in total, of which *N*_*E*_ are excitatory and *N*_*I*_ inhibitory, this constraint is at the origin of the non-accessible areas (white regions).

In the same way as in Fig. 4b, these experiments have been reproduced several times with randomised initial conditions and stimulation sequences. Again thanks to the Kuramoto-Daido order parameters, we were able to quantify over different realizations the number of clusters in the network during the each phase of the protocol according to the number of inputs learned and inhibitory neurons. As before, these analyses gave negligible variations confirming the robustness of the results. In practice, when the networks are randomly initialized, it is rather difficult to reach the optimal configuration representing the upper limit in which each inhibitory neuron controls for one of the *M* clusters. However, we show that by preparing the weight matrices as explained before and by considering a network with 50% of excitatory and inhibitory neurons it is possible even to obtain a “splay state”^84^, characterized by *N/*2 clusters, each formed by a pair of excitatory and inhibitory neurons, spiking one after the other (see Supplementary Figure 1).

Another important aspect to consider is that the time interval (the phase shift) between the population bursts associated to each single cluster reduces with the number of clusters present in the network. Therefore, since the time (phase) potentiation window of the plasticity function Λ(Δ*θ*), shown in Fig. 1, has a finite duration (width) this can induce correlations among clusters characterized by time (phase) shifts smaller than the duration (width) of such a window. These correlations then lead to a merging of initially independent clusters into a single one on the long run. This phenomenon is clearly visible in Supplementary Figure 1, where the “splay state” merges into few clusters over the long term. Therefore, the time (phase) potentiation window should be carefully selected to be sufficiently narrow to reinforce neurons in phase, but also sufficiently broad to allow for some tolerance to the exact phase matching due to noise and variability. To summarize, we can safely affirm that there is a time difference (phase-shift) limit below which nearby clusters cannot remain dynamically independent. At the same time, we should keep in mind that the tolerance of the adaptation process is also a parameter affecting the stabilization of the clusters in the long term.

### Neurons encoding for multiple stimuli

So far we have only considered cases in which external inputs were spatially segregated, meaning that each input targets a separate group of neurons. We now consider the possibility that a sub-group of neurons can encode two stimuli at the same time, thus exhibiting a simple form of “mixed selectivity” often observed in neurons in the prefrontal cortex of primates during the performance of memory related tasks^85,86^. In particular, we study the case in which a sub-group of neurons responds to two distinct stimuli. For illustration we start by considering two stimuli presenting an overlap over eight neurons, see protocol in Fig. 7a. To facilitate the formation of the connections we now strictly alternate the areas stimulated and keep a short resting period between each stimulation.

**Figure 7.**
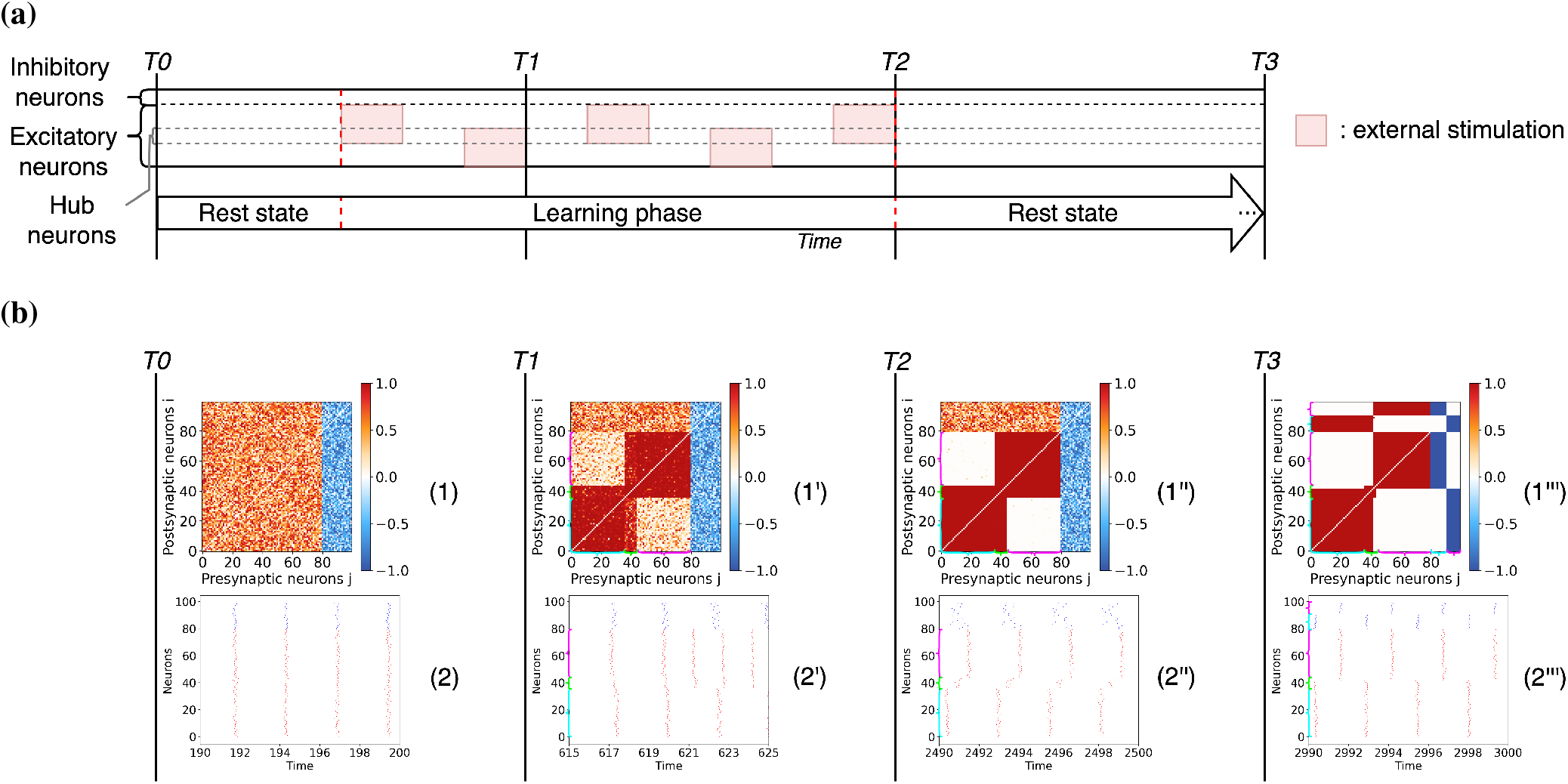
Entrainment of a networks of excitatory and inhibitory *θ*-neurons with two overlapping stimuli. (a) Schema of the experiment protocol showing that the two presented stimuli involve 8 shared neurons. (b) The results are given at different instants of the simulation. The time labels and the graphs have the same significance and content as in Fig. 3. Note that the inhibitory neurons and the putative hub neurons are sorted by phases at time T3 for visualization purposes. The cyan, magenta and green brackets represent clusters 1, 2 and the hubs respectively when they are visible in weight matrices and raster plots.

The results depicted in Fig. 7b are similar to the observations reported before with the formation of two clusters due to the adaptation to the stimuli (see matrix (1”)). The only difference is that in this case a few neurons – those who receive overlapping inputs – become structural hubs. During the consolidation stage, these putative hub neurons associate with one of the two clusters decoupling from the other one but tend to remain connected with the other hubs as shown in matrix (1”‘). Regarding the spiking dynamics, at the end of the training period, the hubs behave as a third independent cluster, by firing at an intermediate time between the two clusters, see raster plot (2”). Although at the end of the consolidation stage they finally become synchronized with one of the two clusters, see raster plot (2”‘). Therefore, this may question on the possibility of exhibiting persistent mixed selectivity with such a model even if the hubs still present structural connections among them. However when increasing the size of the overlap, the two clusters are more likely to become synchronized with each other. This consequently raises the question of what is the largest size of the overlap that allows the two clusters to remain distinct at the end.

To address this question, we generalize the protocol by increasing the number of hubs *N*_*H*_ (neurons that receive both inputs) and the number of inhibitory neurons *N*_*I*_ in the network. For each combination of overlap and number of inhibitory neurons, we observe whether the final network still exhibits two spatially segregated clusters, or if they merge into an all-to-all connected network. The results of this analysis are reported in Fig. 6b: the orange (purple) region refers to the stability (instability) region of the two clusters. As a first constraint observed, the number of excitatory neurons in each cluster cannot be smaller than the number of hubs, i.e. of the number of excitatory neurons shared with the other cluster. Otherwise the hubs would become the dominant cluster leading to synchrony between all excitatory neurons. Furthermore, the number of inhibitory neurons required to maintain the two clusters segregated should be at least twice the number of hubs (*N*_*I*_ ≥ 2 *N*_*H*_). The reason is that to avoid global synchrony, each module needs to compensate for the excitatory links shared with the other cluster via a similar amount of inhibitory links. Therefore the minimal number of inhibitory neurons required to stabilize the two clusters with *N*_*H*_ overlapping neurons is *N*_*I*_ = 2*N*_*H*_ + 1, where an extra inhibitory neuron is needed to maintain the repulsion between the two excitatory modules as previously observed in Fig. 6a. This argument can be generalized to *M* clusters, giving rise to the following rule *N*_*I*_ = (*N*_*H*_ + 1)* *M* −1, where the *N*_*H*_ hubs encode for all the *M* stimuli. However, for sufficiently large *N*_*I*_, namely 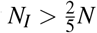, due to the first constraint discussed above, the minimal number of inhibitory neurons needed to observe the two clusters becomes *N*_*I*_ = *N* − 3*N*_*H*_. For an arbitrary number of clusters *M*, this rule generalizes as *N*_*I*_ = *N* − (*M* + 1) * *N*_*H*_. This limit is further confirmed by the analysis reported in Supplementary Figure 5, where we repeat the experiment shown Fig. 7 by reducing the number of inhibitory neurons from *N*_*I*_ = 20 to *N*_*I*_ = 16. In this case, since the minimal number of required inhibitory neurons should be 17, to maintain 2 clusters with 8 hubs, the two clusters indeed merge into an unique one.

Similarly to Fig. 4b, these experiments have been reproduced several times with randomised initial conditions. Here the Kuramoto-Daido order parameters allow to quantify over different realizations if the two clusters remain stable in the network during and after the learning depending on the number of hubs and inhibitory neurons. Again, these analyses gave negligible variations confirming the robustness of the results.

### Memory storage and recall are controlled by the inhibitory neurons

In this last experiment, we start from previous results showing that the connections between excitatory neurons are quite volatile in the absence of learning and that inhibitory connections control for the long-term storage of memories in the cortex^36^. We consider the scenario in which the excitatory clusters formed in the first experiment of Fig. 3 have disappeared due to the volatility of the excitatory connections (e.g. due to new stimulation patterns), while the connections involving inhibitory neurons are maintained since they are more stable. Specifically, we select random connections between the excitatory neurons, while the connectivity involving the inhibitory neurons conserves the values induced by the original learning process. Then we examine the response of the network with this altered connectivity matrix to a brief recall of the stored memory patterns. The schema summarizing the stimulation protocol and the results of this experiment are shown in Figs. 8a and 8b.

**Figure 8.**
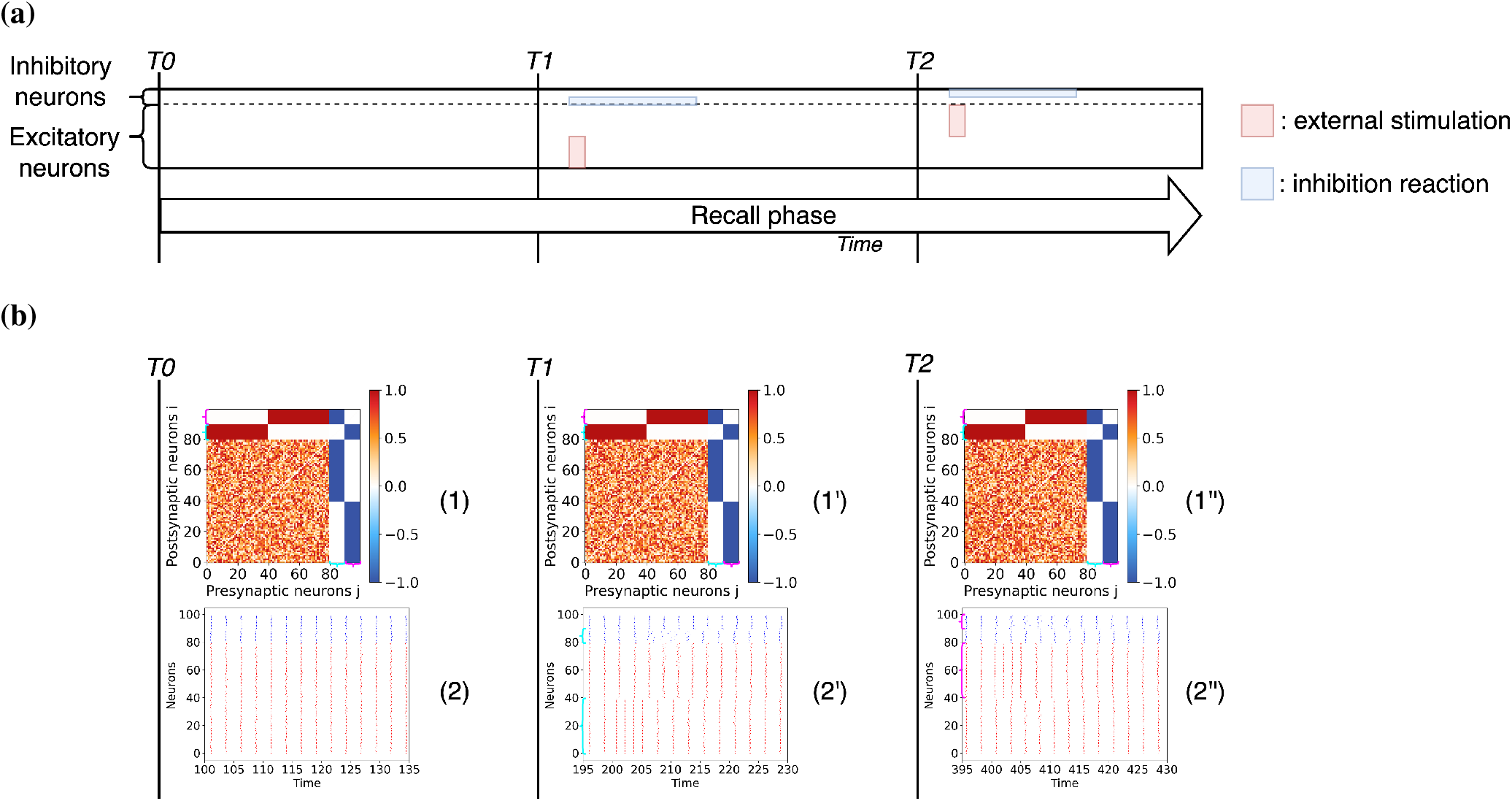
Memory recall experiment for a network of excitatory and inhibitory *θ*-neurons. (a) Schema of the pattern recall experiment. (b) The results are given at different instants of the simulation. The time T0 corresponds to the beginning of the simulation before the stimulations, after a transient period *t*_*t*_ = 100 has been discarded. Time T1 corresponds to the stimulation of the first pattern (from time *t* = 200 to time *t* = 205) followed by the recall of this one. Time T2 corresponds to the stimulation of the second pattern (from time *t* = 400 to time *t* = 405) followed by the recall of this one. The graphs have the same significance as in Fig. 3. The cyan and magenta brackets represent clusters 1 and 2 respectively when they are visible in weight matrices and raster plots.

First of all we observe that when no stimulation is applied to the system, the network is in a relatively synchronized state as shown by the raster plot (2) at time T0. Therefore, the information stored in the network, corresponding to two segregated memory patterns, is not evident from the temporal evolution of the neurons. However, when one of two patterns is presented at times T1 and T2 this induces an initial increase in the spiking rate of the stimulated neurons and shortly after, the desynchronization of the inhibitory neurons associated to the stimulus from the rest of the population for a brief time interval (see raster plots (2’) and (2”)). This leads to a transient re-emergence of the stored memory pattern, reflecting an activity that is almost in anti-phase with the activity of the excitatory neurons associated to other patterns. These results show that for a specific pattern to be recognized it is sufficient that memories are stored in the connections associated with the inhibitory neurons.

## Summary and Discussion

The neural networks of animals, from the simple nervous system of the worm *Caenorhabditis elegans* to the mammalian brain, display modular architectures at different scales of organization^23–28,87–90^. In this paper we have investigated the formation and the consolidation of neural assemblies as driven by the entrainment to different inputs in networks of oscillatory *θ*-neurons. Previous analyses have shown that learning and adaptation could generate modular networks^16,29,91,92^, but as it is common practice in artificial neural networks, those studies overlooked some fundamental biological constraints. Here, we have shown that satisfying Dale’s principle – i.e. the distinction between excitatory and inhibitory neurons – is crucial for the consolidation of stimulus-driven neural assemblies. Furthermore, at variance with other popular models of learning where the synaptic weights and the neural activity are frozen once the training phase is finalized, we allowed for a spontaneous activity of the adaptive network even after the training was finished, thus mimicking a more biologically realistic scenario. We have found that during this post-training phase the learned memories are consolidated, provided the network is made of both excitatory and inhibitory neurons. Indeed, if the network contains only excitatory neurons, then the learned memories are lost during to the post-learning phase. Moreover, if the network contains excitatory and inhibitory synapses, but Dale’s principle is violated, the network ends in a state in which the emerged synaptic connectivity does not correspond to the encoded stimuli. Furthermore, we have shown that the number of inhibitory neurons controls for the memory capacity of the network.

### Create, maintain and consolidate memories

We have seen that the creation of modular structures is driven by an adaptation mechanism in which excitatory neurons receiving the same input reinforce their connections while the connections between uncorrelated neurons are essentially removed. The learning process allows these connected neurons to maintain a certain degree of synchrony among them while still exhibiting some variability in their dynamics due to the external noise and the heterogeneity in their parameters. The structure emerging in the network due to the learning phase resembles the modular and hierarchical organisation of the brain where each modules represent a sensory modality^23,24,26,87,93^

Regarding the maintenance of the generated structural modules, we observed that the inhibitory neurons play a fundamental role in preventing a global resynchronization of the whole network after the learning phase. Each inhibitory neuron becomes associated with a particular group of excitatory neurons: it receives excitation from this group while it inhibits the rest of the excitatory neurons as illustrated in Fig. 4a. Therefore, each cluster exhibits an independent dynamics, preventing a long-term forgetting of the structural neural assemblies, each coding for a different memory item (stimulus). At variance to what has been reported in previous studies on excitatory-inhibitory networks with chemical synapses where sufficiently strong inhibitory connections and strong excitatory-inhibitory coupling favour the synchronization in the network^94,95^, in our system the inhibitory neurons are also synchronized by the excitatory ones, but their activity tends to maintain the excitatory clusters desynchronized. This difference can be understood by noticing that in our case the couplings among the neurons are essentially electrical gap junctions, which for moderate strength and in absence of delay are known to promote in-phase (anti-phase) dynamics among excitatory (inhibitory) neurons^96^ similarly to what is observable for phase oscillators.

We have carried out several additional experiments to validate the robustness of the results and to better understand the mechanisms at the basis of memory storage and consolidation. On the one hand, we have shown that stored memories can be retrieved with a brief stimulation recall even if the excitatory assemblies associated to the presented stimuli are no longer present in the connectivity, but are preserved only in the connections involving the inhibitory neurons, as shown in Fig. 8b. In the light of these results, one could speculate that the main purpose of the connections between excitatory neurons is to store short-term memories while the links associated with the inhibitory neurons – which have been reinforced during the consolidation post-training phase – correspond to long-term memory storage. As long as inhibitory neurons preserve their connections, short-term memories could be erased to process and learn new information. This result resembles the one found in^41^ for excitatory-inhibitory networks with inhibitory plasticity.

On the other hand, we have shown the possibility of consolidating structures despite the learning did not involve all excitatory neurons. In particular, we have studied the case in which partially trained areas were not completely formed due to partial random stimulations and we have analyzed the case where a sub-group of neurons remains untrained (unstimulated), see Supplementary Figures 2 and 3. From these analyses, we can conclude that even if the system partially learns (but at a sufficient degree) the given stimulation patterns, then the connections will be anyway reinforced during the post-learning phase. Here, an analogy could be drawn with memory consolidation during sleep^83,97^, when the memories of the daytime experiences are recalled for their reinforcement. Finally, we have shown that the reported results are general by replicating the experiments with two other oscillator models (i.e. Kuramoto and Stuart-Landau model) as shown in Supplementary Figure 6.

### Memory capacity of the network

In Fig. 6a it is shown that the number of inhibitory neurons is linked to the number of different neural assemblies that may keep an independent (not synchronised) dynamics and consequently be consolidated. If we consider each cluster as a stored memory item, we can easily link the number of inhibitory neurons to the capacity of a network to learn and store information. Therefore, we can safely affirm that the number of inhibitory neurons is clearly related to the memory capacity of the network^37^. In particular we have shown that for non-overlapping memories the maximal capacity is proportional to the number of inhibitory neurons, thus we expect that the maximal storage capacity will grow as ≃ 0.20*N* by assuming that 20 % of the neurons are inhibitory as observed in the cortex. This capacity is definitely larger than that of the Hopfield model in which the nature of excitatory and inhibitory neurons is not preserved and whose memory capacity can grow at most as ≃ 0.14*N*^19^. Furthermore, we have shown in Fig. 6b that the number of inhibitory neurons controls also the maximal number of neurons which can code different items at the same time, i.e. neurons exhibiting a simple form of “mixed selectivity”^85^. These neurons could be seen also as hub neurons allowing for connections between the stored clusters and facilitating the ability to potentially transmit and integrate information^26,93,98,99^. Once again we have shown that the amount of inhibition can be related with the cognitive capability of the network.

### Limitations and outlook

Despite the efforts performed in the present work to fulfill several biologically plausible constrains, some aspects of the model could be further improved. For example, a more realistic scenario could replace the *θ*-neuron model here employed by spiking neurons with STDP plasticity. The learning rule considered here corresponds to the basic paradigm of an Hebbian rule implying that “*cells that fire together, wire together*”^100^ and depends on the difference of the phases associated to each neuron in a continuous manner. By employing this rule, since synaptic potentiation occurs only when two phases match within a quite narrow window, we have shown that a lack of precision in the measurements of the phase difference can have a real impact. In particular, we have shown how this lack of precision can limit the number of memory items that can be effectively stored in the long term, in Supplementary Figure 1. Another crucial element of the model is the presence of two time scales associated to the evolution of the synaptic weights, a faster adaptation during the learning phase and a slower plasticity evolution during rest. While this choice is justified by several indications that learning and consolidation processes may occur simultaneously or sequentially in the brain characterized by different time-scales^64–67,72–75^, it is still unclear how the brain itself solves this problem and avoids the loss of learned memories given the fact that synapses are permanently susceptible to adaptation. Better understanding of the biological mechanisms will allow to define more realistic adaptation rules in the models. Similarly, multi-scale mechanisms are present and necessary in real brain activity, since the sensory information transmitted to the brain are not all learned instantaneously and some reach it passively while others need more time and repetition to be assimilated. In this way, it would be interesting to detect the novelty or the relevance of the inputs in order to adapt the weights at different time-scales, thus opening the door to the development of more flexible systems that are capable of learning more efficiently. Additionally, we foresee that some of the computational experiments here performed could be carried out *in vitro*, for example, by considering neuronal cultures with or without inhibitory neurons and by applying different patterns of localised electrical or opto-genetical stimulations to sub-groups of neurons.

## Data Availability

The datasets used and/or analysed during the current study available from the corresponding author on reasonable request.

## Code Availability

All code for algorithms and analysis associated with the current submission is available at https://github.com/rbergoin/Phase_models_neural_assemblies.

## Acknowledgments

This work was supported (R.B., G.D. and G.Z.) by the European Union’s Horizon 2020 Framework Programme for Research and Innovation under the Specific [Grant Agreement No. 945539 (Human Brain Project SGA3)] and by an EUTOPIA funding [EUTOPIA-PhD-2020-0000000066 - NEUROAI]. A.T. received financial support by the Labex MME-DII [Grant No. ANR-11-LBX-0023-01] (together with M.Q.), and by the ANR Project ERMUNDY [Grant No. ANR-18-CE37-0014] all part of the French program “Investissements d’Avenir”. G.D. is supported by the Spanish national research project [ref. PID2019-105772GB-I00/AEI/10.13039/501100011033] funded by the Spanish Ministry of Science, Innovation, and Universities (MCIU). M.Q. is also partially supported by CNRS through the IPAL lab in Singapore. A.T. thanks G. Mongillo and S. Olmi for useful discussions.

## Author contributions

R.B., G.Z., A.T. and M.Q. conceived the experiments, R.B. conducted the experiments, R.B. analysed the results. R.B. wrote the manuscript. All authors reviewed the manuscript.

