## Supplementary Information for "Inhibitory neurons control the consolidation of neural assemblies via adaptation to selective stimuli"

#### Alternative Stimulation Protocols

In this section we present results relative to additional stimulation protocols which consist of variations of the initial protocols depicted in the section *Methods* of the main text. The dynamics are simulated using the $\theta$-model as in the main text.

**Emergence of splay states** The idea of this experiment is to start from the connectivity matrix shown in panel (1) of the Supplementary Figure 1 where all the excitatory neurons are decoupled one from another, but where each of them has a single post-synaptic connection to an inhibitory neuron. The latter consequently inhibits all neurons except the one with which it is associated. As evident from the raster plot (2) displayed at time T0 of the Supplementary Figure 1, all pairs of excitatory-inhibitory neurons form pseudo-clusters that are desynchronized from the other pairs forming a splay state. Certainly, this represents an ideal limiting case since we artificially create the structure without recurring to any learning process. Nevertheless, it represents the extremal case in which a maximum number of possible clusters can be generated, confirming that one inhibitory neuron per cluster is theoretically sufficient to maintain each pair of neurons desynchronized from the others.

However, after a long period of spontaneous activity and consolidation, this structure and this particular dynamics are not maintained. Indeed as explained in sub-section *Learning multiple clusters* of the main text, the phase (time) potentiation window of the plasticity function determines the interval within which the neurons are considered to be correlated. The clusters forming the splay state spike at very close (though strictly different) times, however the plasticity forces the temporally close clusters to merge over the long term, as evident in the weight matrix (1') obtained at long time after consolidation (T1). Thus, the network reaches a stable state with about 5 clusters spiking at distinct times (see raster plot (2')).

**Two memory patterns with randomly stimulated neurons** In the previous experiments, when a group of neurons was stimulated, all the neurons of the group received the same stimulation input. In this experiment, only a certain percentage of the neurons within the group receive the stimuli. The neurons are randomly selected with a probability of 50%. The protocol and the results corresponding to this experiment are depicted in Supplementary Figures. 2a and 2b.

The first difference observed here is that in the connectivity matrix (1") obtained after the learning (time T2), the clusters are less pronounced due to the random nature of the activation of the neurons that constitute each of them. Despite this, the two groups can be still distinguished, firing almost together at different times (raster plots (2")). Afterwards, the reinforcement phase (time T3) allows to obtain the same connectivity and dynamical evolution as in the original protocol. The interest of this protocol is to demonstrate the possibility of creating memory patterns with more realistic input signals. Indeed in the brain, not all the neurons in an area are stimulated at the same time.

**Three groups of excitatory neurons, with only two subject to learning** The idea of this experiment is to reproduce the same original protocol but this time keeping a group of excitatory neurons unstimulated and therefore not subject to a strong

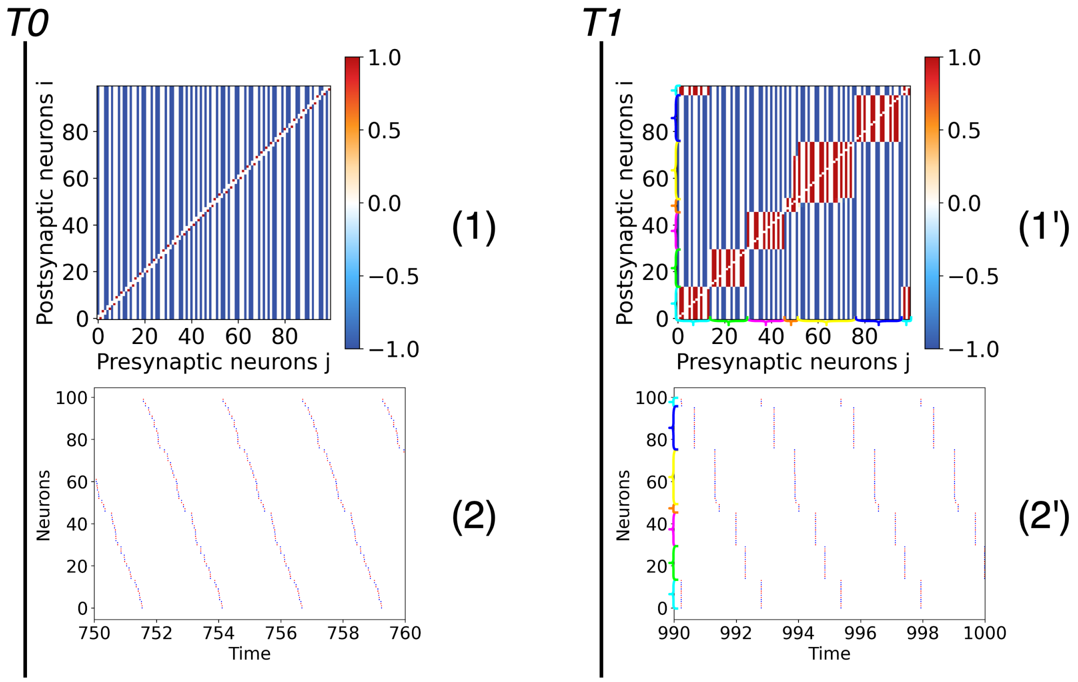


**Supplementary Figure 1.** Splay states in the $\theta$-model. Results of the experiment where the connectivity matrix is initialized to exhibit $N/2$ clusters corresponding to a splay state. The time T0 corresponds to the beginning of a resting phase in absence of any stimulations, after a short transient period $t_t = 750$ has been discarded. The time T1 corresponds to the end of a long period of spontaneous activity during which synaptic weights are consolidated. Panels labeled (1) and (1') represent the weight matrices at times T0 and T1, respectively: the color denotes if the connection is excitatory (red), inhibitory (blue) or absent (white). Panels (2) and (2') are raster plots at times T0 and T1, displaying the firing times of excitatory (red dots) and inhibitory (blue dots) neurons. Note that in both figures, for clarity the neurons are sorted by phases. Also to better visualize the splay state, a homogeneous system without noise is considered in this experiment. The cyan, green, magenta, orange, yellow and dark blue brackets represent clusters 1, 2, 3, 4, 5 and 6 respectively in weight matrices and raster plots after the consolidation phase.

adaptation. See the protocol of this experiment and its results in Supplementary Figures. 3a and 3b.

The first observation after the learning phase (time T2) is that the two structural clusters created, connectivity matrix (1"), are also visible in the raster plots as two groups of neurons firing in anti-phase (2"). However, the group of unstimulated excitatory neurons fires quite irregularly at this time as the inhibitory neurons. This similarity with the inhibitory group is also observable after the consolidation process (time T3) where the unstimulated excitatory neurons associate with one of the two excitatory clusters and its associated inhibitory neurons. Thus, we can clearly see in matrix (1"') that both the untrained excitatory neurons and the inhibitory population are splitted into two structural clusters, each of them associated to one of the two memory patterns making them grow. Depending on the initial conditions it can happen that the two groups are not equally populated and in extreme cases the unstimulated excitatory neurons can associate also to only one of the two memory patterns.

Note that this phenomenon is only possible when the original memory patterns involve a sufficient number of neurons with respect to the unstimulated group. When the number of the latter prevails, the two stimulated groups tend to synchronize and only one pattern emerges in the end. As discussed in the multi-cluster learning, the number of inhibitory neurons associated to each excitatory cluster has also a role in determining which is the minimal number of excitatory neurons in each memory pattern that can remain separated on the long run.

**Three memory patterns learned, two maintained** This experiment is analogous to the experiment reported in Fig. 5 in the sub-section *Learning multiple clusters* of the main text. In particular, three different stimuli are presented to the excitatory neurons as in the main text, but now only one inhibitory neuron is present in the network. The results are reported in the Supplementary Figure 4.

At the steps T0 and T1, we logically obtain similar results as in the main text. However at time T2 after the learning phase, although the three clusters are well formed as before (see connectivity matrix (1") in Supplementary Figure 4b), two of them share a close dynamics, i.e. almost synchronized (as shown in in raster plot (2") in Supplementary Figure 4b). The direct

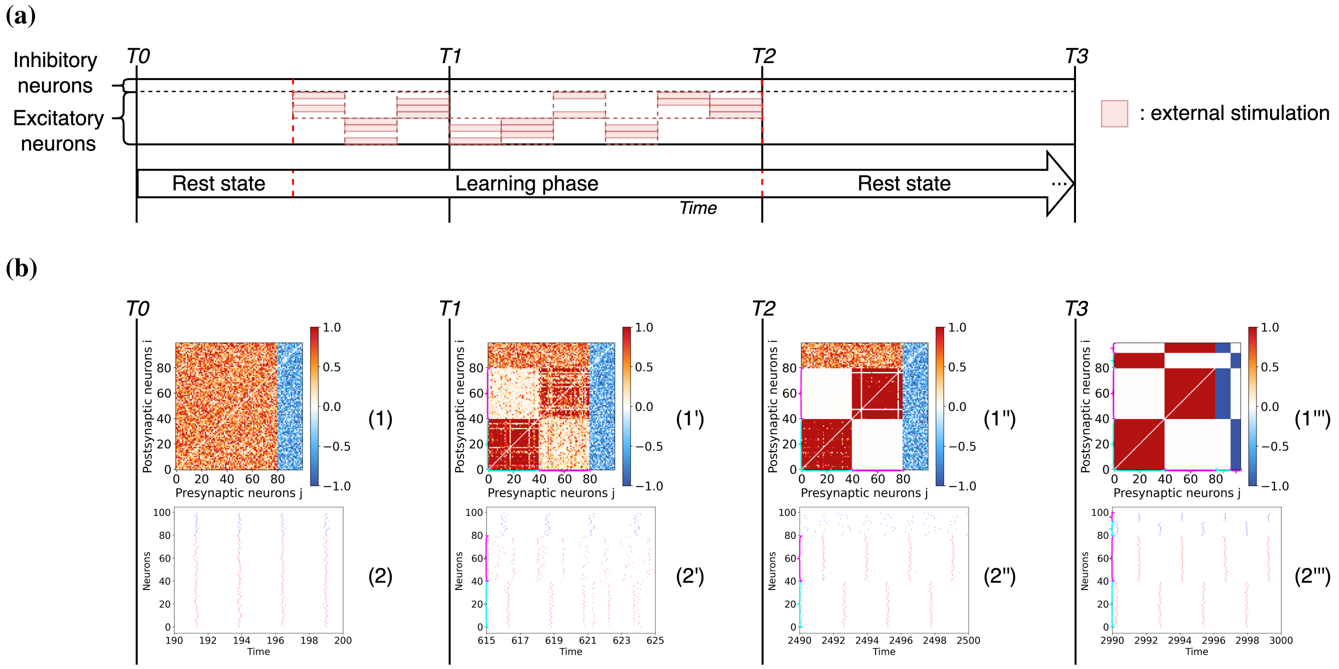



**Supplementary Figure 2.** Entrainment of a networks of excitatory and inhibitory $\theta$-neurons with two stimuli applied to random subsets of neurons within each group. (a) Schema of the experiment consisting of the stimulation of randomly chosen excitatory neurons of each of the two groups every stimulation period. (b) The results are given at different instants of the simulation. The time labels and the graphs have the same significance and content as in the main text. Note that the inhibitory neurons are sorted by phases at time T3 for visualization purposes. The cyan and magenta brackets represent clusters 1 and 2 respectively when they are visible in weight matrices and raster plots.

consequence of this is that in the long term (time T3), these two structural modules merge in only one (panel (1"')) and they are now completely synchronized (panel (2"')). In other words, one of the three learned items is forgotten and finally only two are memorized. This experiment confirms the previously established limit by showing that one inhibitory neuron is not sufficient to maintain three memories ($N_{inhibitory} = 1 \not\geq M - 1 = 2$ with $M = 3$ clusters). However, it also shows that a single inhibitory neuron is sufficient for maintaining two memories ($N_{inhibitory} = 1 \geq M - 1 = 1$ with $M = 2$ clusters).

**Unstable hub neurons** In this last alternative stimulation protocol, we reproduce the same experiment shown in Fig. 7 in the sub-section *Neurons encoding for multiple stimuli* in the main text. As in the original case, we allow the two stimuli to share eight neurons among them but this time there are only sixteen inhibitory neurons in the network. The results of the experiment are reported in the Supplementary Figure 5.

During the steps T0 and T1, as expected we obtain similar results to the one reported in the main text. However at time T2 after the learning phase, although the two clusters and the "hub neurons" are well formed as in the main text (see connectivity matrix (1")) in the Supplementary Figure 5b), now the two clusters almost synchronize (see raster plot (2") in the Supplementary Figure 5b). Consequently over the long post-training period (time T3), the clusters merge into one (panel (1"')) and they become completely synchronized (panel (2"')). Therefore, in addition to being unstable over the long term, these hub neurons make the modules unmaintainable. As speculated in the main text, the excitatory connections between modules that are mediated by the hubs must be compensated by an equivalent amount of inhibition. This corroborates the previously established limit by showing that sixteen inhibitory neurons are not sufficient to maintain eight hubs ($N_{inhibitory} = 16 \not\geq 2 * M + 1 = 17$ with $M = 8$ hubs).

### Networks of oscillators

Throughout this study, single units have been simulated considering the $\theta$-neuron model. We now reproduce the results of Fig. 3 in the main text considering networks of phase oscillators (the Kuramoto model) and of oscillators characterized by their phase and amplitude (the Stuart-Landau model). The adaptation of the synaptic weights will be governed by the same rule employed for the $\theta$ model and reported in the *Methods* section.

**The Kuramoto model** The Kuramoto model for coupled phase oscillators has the advantage of being a relatively simple model able to capture different types of dynamics ranging from asynchronous to partially synchronous states[1–3]. Some

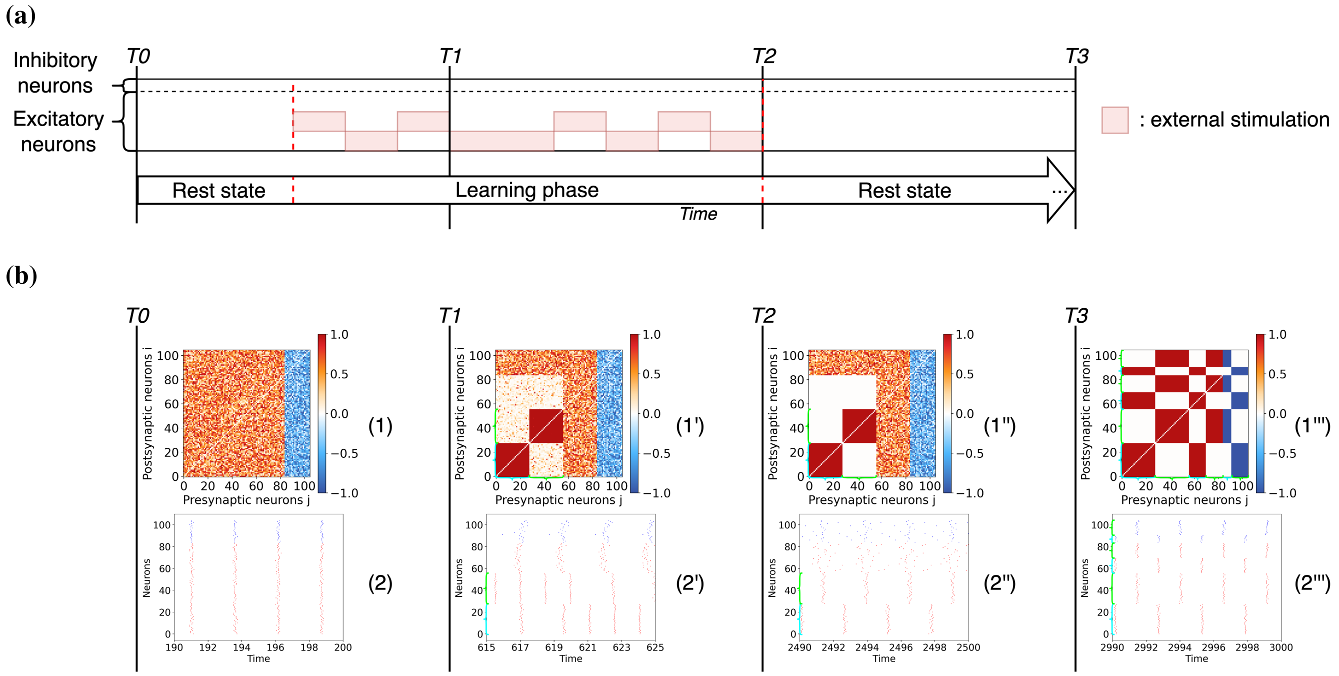


**Supplementary Figure 3.** Entrainment of a network of excitatory and inhibitory $\theta$-neurons with two non-overlapping stimuli and a group of free excitatory neurons. (a) Stimulation protocol with two stimulated groups of excitatory neurons and another group of unstimulated neurons. (b) The results are given at different instants of the simulation. The time labels and the graphs have the same significance and content as in the main text. Note that the inhibitory neurons and the initially untrained excitatory neurons are sorted by phases at time T3 for visualization purposes. The cyan and green brackets represent clusters 1 and 2 respectively when they are visible in weight matrices and raster plots.

studies have shown the possibility of adding an adaptation mechanism to the coupling leading to more complex dynamical behaviours[4–7]. Therefore, it is important to compare our findings for a classical neuronal model with the paradigmatic Kuramoto model, where the evolution of the phase $\theta_i$ of an oscillator is described by:

$$\frac{d\theta_i}{dt} = \omega_i + \frac{g}{N}\left(\sum_{j=1}^{N} \kappa_{ij}\sin(\theta_j - \theta_i)\right) + I_i(t) + \xi_i(t) \quad . \tag{1}$$

The natural frequency of oscillator $i$ is denoted by $\omega_i$, $g$ represents the global coupling strength, while $\kappa_{ij}$ is the relative directed coupling from the oscillator $j$ towards the oscillator $i$. In this context, $I_i(t)$ represents an external driving term and $\xi_i(t)$ represents an additive Gaussian noise.

**The Stuart-Landau network model** One of the limitations of the Kuramoto model is the unrealistic nature of its dynamics compared to biological oscillators, in particular the fact that the evolution of the amplitudes is neglected. The normal form for a non-linear oscillator in the proximity of a Hopf bifurcation is represented by the Stuart-Landau model which describes not only the phase variations but also the amplitude variations of the oscillator. The dynamics of this oscillator is given in terms of a complex variable $z_i = \rho_i e^{\hat{\imath}\theta_i} = x_i + \hat{\imath} y_i$ where $\rho_i$ represents the amplitude of the $i$th unit, $\theta_i \in [-\pi, \pi[$ its phase and $\hat{\imath}^2 = -1$. The evolution of the $i$-th oscillator in a network is described by Deco et al.[8]:

$$\frac{dz_i}{dt} = z_i\left[\alpha_i + \hat{\imath}(\omega_i + I_i(t)) - |z_i^2|\right] + \frac{g}{N}\left(\sum_{j=1}^{N} \kappa_{ij} z_j\right) + \xi_i(t) \tag{2}$$

Most of the quantities appearing in Eq. (2) have been already defined in Eq. (1). The only new element is the real parameter $\alpha_i$ which governs the type of dynamics of the system: $\alpha_i < 0$ leads to a stable fixed point and $\alpha_i > 0$ to a stable limit cycle oscillation[8]. The Gaussian noise $\xi_i(t)$ is here considered to be complex.

In order to guarantee a similar firing rate at rest (where the spike is defined like in the $\theta$-neuron model) and a stimulation of the same order, we have adapted the specific parameters of these two models as indicated in table 1. Parameters not indicated in the table are considered identical to the ones employed for the $\theta$-neuron model.

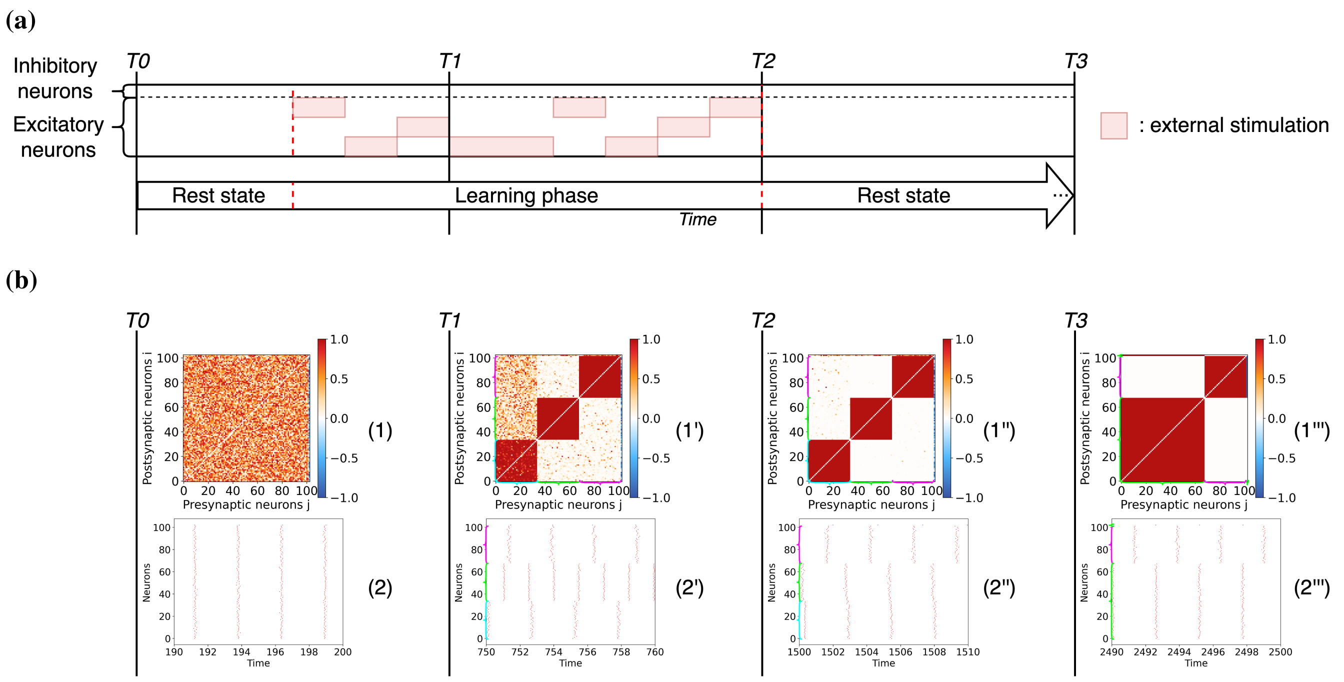


**Supplementary Figure 4.** Entrainment of a network of excitatory and inhibitory $\theta$-neurons with three stimuli in presence of only one inhibitory neuron. (a) Stimulation protocol of the excitatory neurons via three different non-overlapping stimuli. (b) Entrainment results at different instants of the simulation. The time labels and the graphs have the same significance and content as in the main text. The cyan, green and magenta brackets represent clusters 1, 2 and 3 respectively when they are visible in weight matrices and raster plots.

| Parameters | Values |
|---|---|
| $\omega_i$ | $\mathcal{N}(2.45, 0.02)$ |
| $\alpha$ | $\mathcal{N}(0.0, 0.2)$ |
| $g$ | 2 |
| $I(t)$ | $\{0, 3.46\}$ |
| $\xi(t)$ | $\mathcal{N}(0.0, 0.2)$ |

**Supplementary Table 1.** Parameters for the Kuramoto and Stuart-Landau oscillator networks

Following the same stimulation protocol as in Fig. 2 of the main text (see Supplementary Figure. 6a), the corresponding results for the $\theta$-model, the Kuramoto and the Stuart-Landau models are reported in Supplementary Figures. 6b – 6d. Essentially, the same results are obtained for the three models as seen in the emerging weight matrices after learning (1”) and reinforcement (1”’), and in the corresponding spiking behaviour (raster plot (2”’)). The Kuramoto and the Stuart-Landau oscillators seem just a little noisier than the $\theta$-neurons. We have obtained with the Kuramoto and Stuart-Landau network models quite similar results also for all the other protocols and numerical experiments performed in the main text for the $\theta$-neuron. These results prove our findings to be quite general and not limited to a specific model of oscillator.

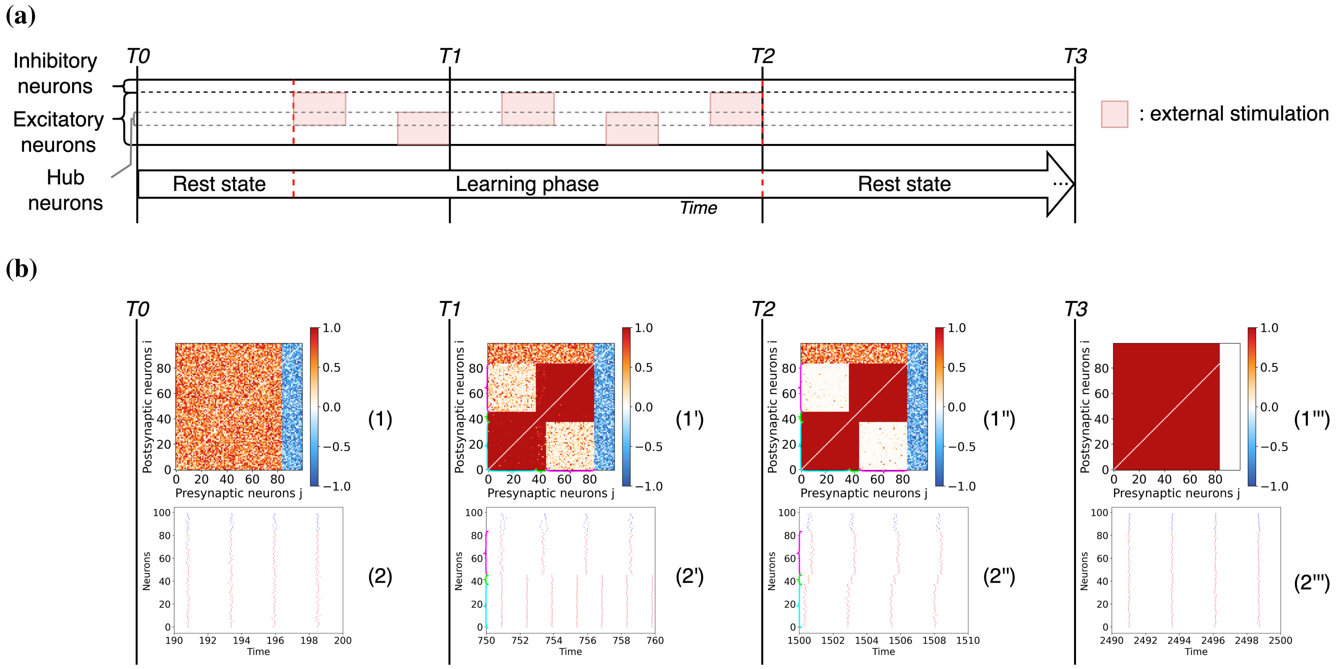


**Supplementary Figure 5.** Entrainment of a network of excitatory and inhibitory $\theta$-neurons with two overlapping stimuli and 16 inhibitory neurons. (a) Schema of the experimental protocol showing that the two presented stimuli involve 8 shared neurons. (b) The results are given at different instants of the simulation. The time labels and the graphs have the same significance and content as in the main text. The cyan, magenta and green brackets represent clusters 1, 2 and the hubs respectively when they are visible in weight matrices and raster plots.

**Supplementary Figure 6.** Entrainment of networks of 80% excitatory and 20% inhibitory neurons for three different oscillatory models. (a) Schema of the experiment leading to the emergence of two clusters due to the stimulation of two non-overlapping excitatory neuronal populations. (b) Results for a network of $\theta$-neurons, (c) for a network of Kuramoto phase oscillators and (d) for a network of Stuart-Landau oscillators at different instants of the simulation. The temporal labels and the graphs have the same significance and content as in the main text. Note that the inhibitory neurons are sorted by phases at time T3 for visualization purposes. The cyan and magenta brackets represent clusters 1 and 2 respectively when they are visible in weight matrices and raster plots.

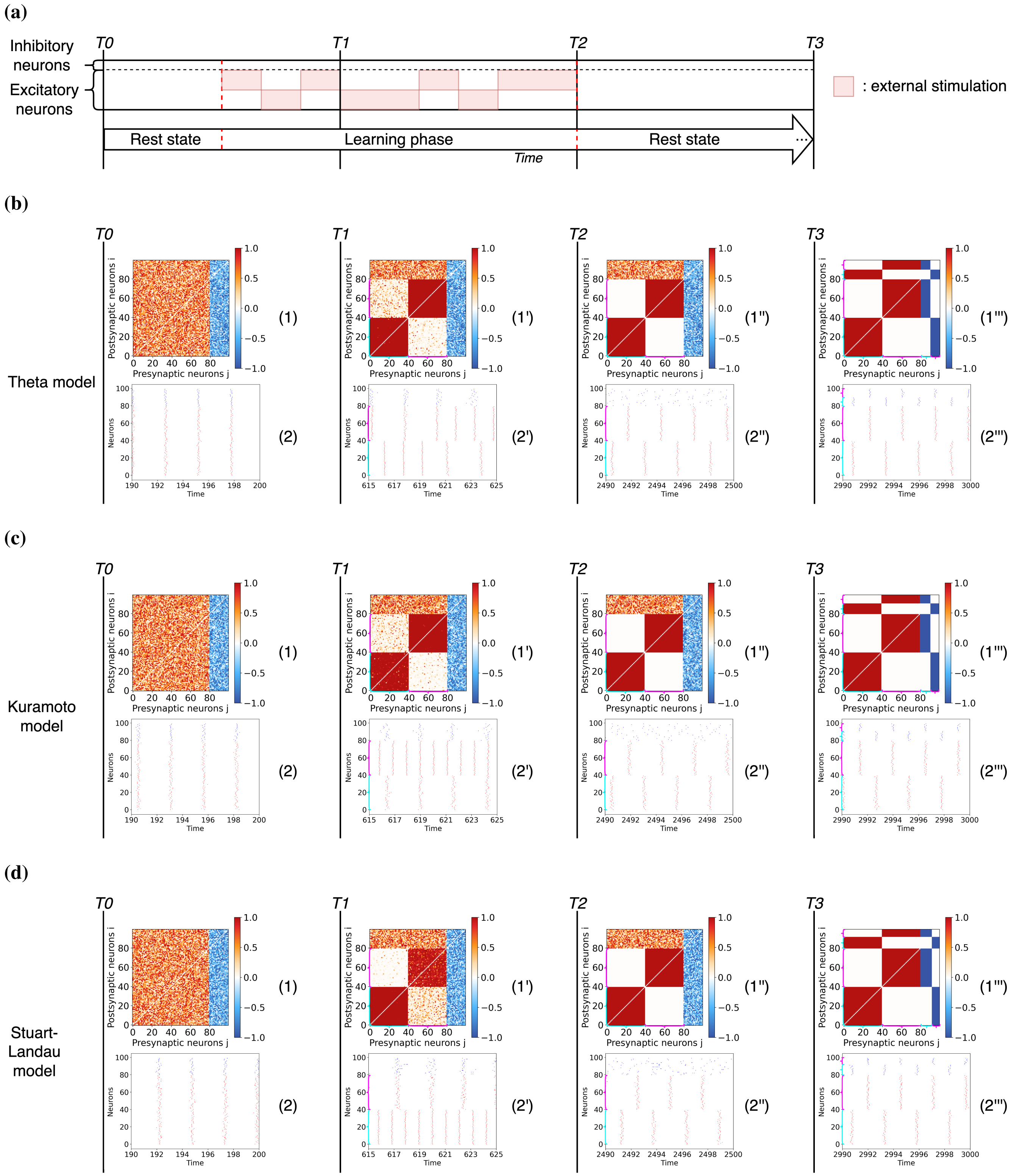
(a)
Inhibitory neurons
Excitatory neurons
T0
T1
T2
T3
: external stimulation
Rest state
Learning phase
Rest state
Time
(b)
Theta model
(c)
Kuramoto model
(d)
Stuart-Landau model
Postsynaptic neurons i
Presynaptic neurons j
Neurons
Time
(1)
(1')
(1'')
(1''')
(2)
(2')
(2'')
(2''')
